# Accurate age prediction from blood using of small set of DNA methylation sites and a cohort-based machine learning algorithm

**DOI:** 10.1101/2023.01.20.524874

**Authors:** Miri Varshavsky, Gil Harari, Benjamin Glaser, Yuval Dor, Ruth Shemer, Tommy Kaplan

## Abstract

Chronological age prediction from DNA methylation sheds light on human aging, indicates poor health and predicts lifespan. Current clocks are mostly based on linear models from hundreds of methylation sites, and are not suitable for sequencing-based data.

We present GP-age, an epigenetic clock for blood, that uses a non-linear cohort-based model of 11,910 blood methylomes. Using 30 CpG sites alone, GP-age outperforms state-of-the-art models, with a median accuracy of ~2 years on held-out blood samples, for both array and sequencing-based data. We show that aging-related changes occur at multiple neighboring CpGs, with far-reaching implications on aging research at the cellular level. By training three independent clocks, we show consistent deviations between predicted and actual age, suggesting individual rates of biological aging.

Overall, we provide a compact yet accurate alternative to array-based clocks for blood, with future applications in longitudinal aging research, forensic profiling, and monitoring epigenetic processes in transplantation medicine and cancer.

**Graphical abstract:** 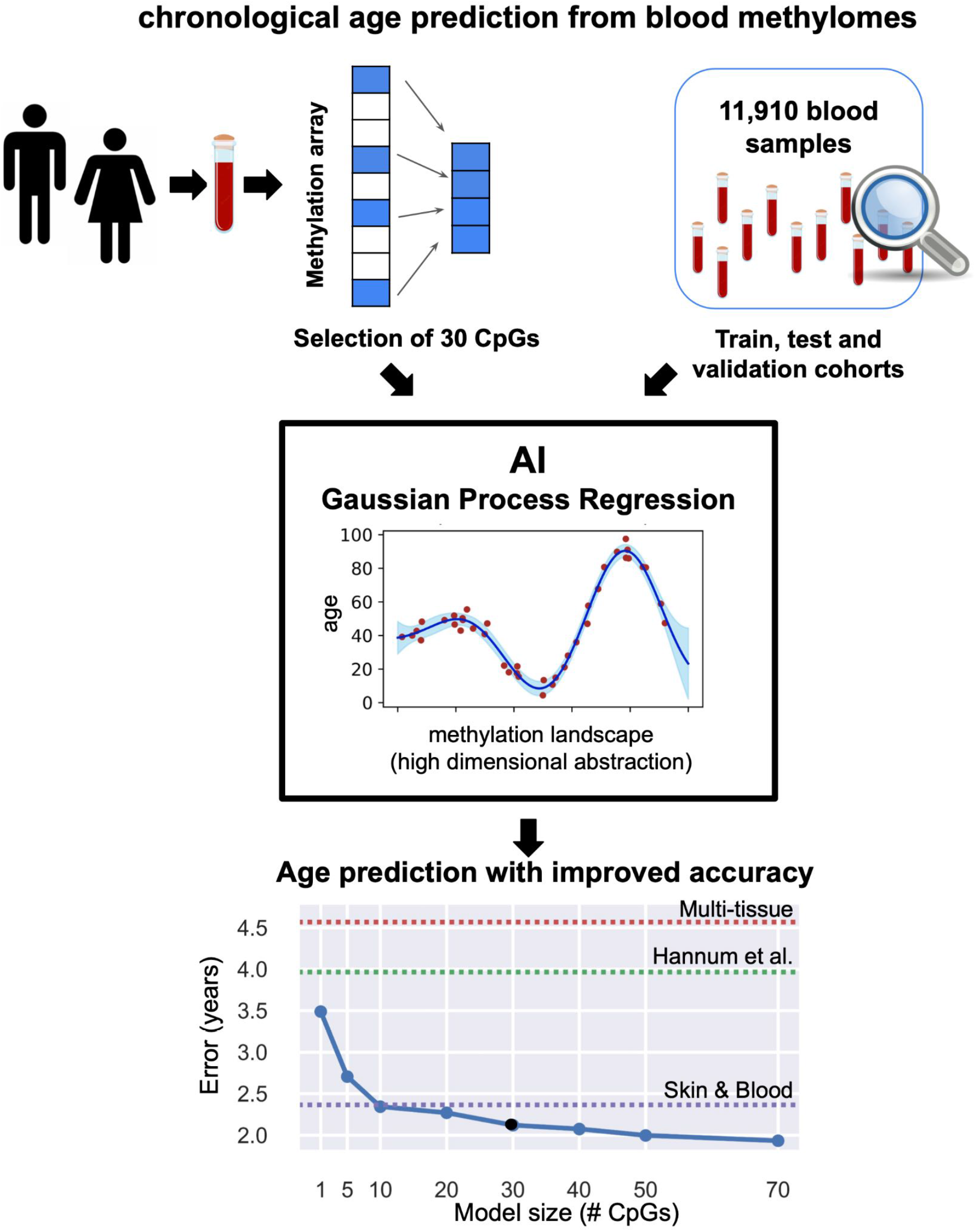

- Machine learning analysis of a large cohort (~12K) of DNA methylomes from blood
- A 30-CpG regression model achieves a 2.1-year median error in predicting age
- Improved accuracy (≥1.75 years) from sequencing data, using neighboring CpGs
- Paves the way for easy and accurate age prediction from blood, using NGS data

**Motivation:** Epigenetic clocks that predict age from DNA methylation are a valuable tool in the research of human aging, with additional applications in forensic profiling, disease monitoring, and lifespan prediction. Most existing epigenetic clocks are based on linear models and require hundreds of methylation sites. Here, we present a compact epigenetic clock for blood, which outperforms state-of-the-art models using only 30 CpG sites. Finally, we demonstrate the applicability of our clock to sequencing-based data, with far reaching implications for a better understanding of epigenetic aging.

## Introduction

Chronological age prediction using DNA methylation emerged from seminal studies by Horvath^1^, and Hannum and colleagues^2^. Both used Illumina DNA methylation arrays, and integrated the methylation levels at a predefined set of 353 or 71 CpG sites (respectively) across the genome in a linear regression model to predict age, yielding impressive predictions. Intriguingly, these models inherently assumed faster changes in DNA methylation levels until the age of 20 years, after which methylation levels were modeled at a constant rate throughout adulthood^1^.

The molecular mechanisms of aging are yet to be fully uncovered, and the factors that drive changes in DNA methylation with age are not well understood. These changes may either be consistent and predictable, such as alterations in DNA methylation as a result of accumulating stress^3^, or spontaneous and unpredictable, such as under-performance of the DNA methylation maintenance system after DNA replication.

Epigenetic clocks are a valuable tool in the research of human aging and the genetic and environmental factors that influence it^4^. In addition to opening a unique window for research opportunities, the prediction of chronological age from methylation data may have applications in multiple fields, such as analysis of DNA samples from crime scenes; indication of poor health and all-cause mortality prediction using “age acceleration” metric, defined as the difference between chronological and predicted age^5–8^; identification of early graft-versus-host disease (GvHD) in organ transplant recipients^9^ and more.

New and improved epigenetic clocks for chronological age prediction have been developed in recent years, using alternative sets of CpG sites. Most clocks were developed as a linear model based on the Illumina BeadChip platform, some using hundreds of CpG sites^10,11^, and others using only a few CpGs^12,13^. Few non-linear models were introduced, including those using neural networks^14,15^. The high cost of the Illumina BeadChip platforms hinders accessible use of such chronological age predictors, and thus several less-accurate clocks trained on small pyrosequencing datasets with few CpG sites were also proposed, mostly for forensic use^16–18^.

In addition to chronological age predictors, the field of biological age predictors has recently gained much attention. These predictors integrate DNA methylation levels with additional clinical biomarkers to predict an individual’s healthspan and lifespan, including their risk for mortality, physical functioning, cognitive performance, and more^8,19,20^. While these models provide a broader view on health and aging, they do not serve the same purpose as chronological age predictors, and are limited to a small number of donors for which multi-omics integrative data were collected, as well as detailed clinical information.

In this work we present GP-age, a non-parametric cohort-based epigenetic clock for chronological age prediction from blood samples, based on a Gaussian Process regression (GPR) model. We collected a large cohort of 11,910 human blood methylomes, measured for healthy individuals at a wide variety of ages, and identified a small set of 30 age-related CpGs. Given a query blood sample, the methylation levels at these sites are compared against the training cohort, and the ages of similar samples are integrated to predict age. As we show, using a small set of 10-30 carefully selected CpGs, GP-age outperforms larger state-of-the-art models. Finally, we demonstrate the applicability of GP-age to studying aging using methylation arrays or sequencing data, providing an accessible and accurate alternative to current clocks and opening new avenues for the study of epigenetic changes at multiple neighboring age-related CpG sites.

## Results

### A dataset of 11,910 blood-derived methylomes of healthy donors across various ages

We assembled a large dataset of publicly-available blood-derived methylomes from 19 genome-wide methylation array studies, obtained from blood samples in a variety of ages and scenarios^2,21–37^.

Overall, our data contains 11,910 blood methylomes, from donors aged 0-103 years (Fig. 1). One of these studies^37^, a large dataset (n=665) that spans a wide range of ages, was selected as a held-out independent validation set, to unbiasedly assess our prediction results. Importantly, this database was not included in the training of previously published epigenetic clocks, allowing a direct and unbiased comparison between GP-age and other state-of-the-art models^1,2,10^.

**Figure 1:**
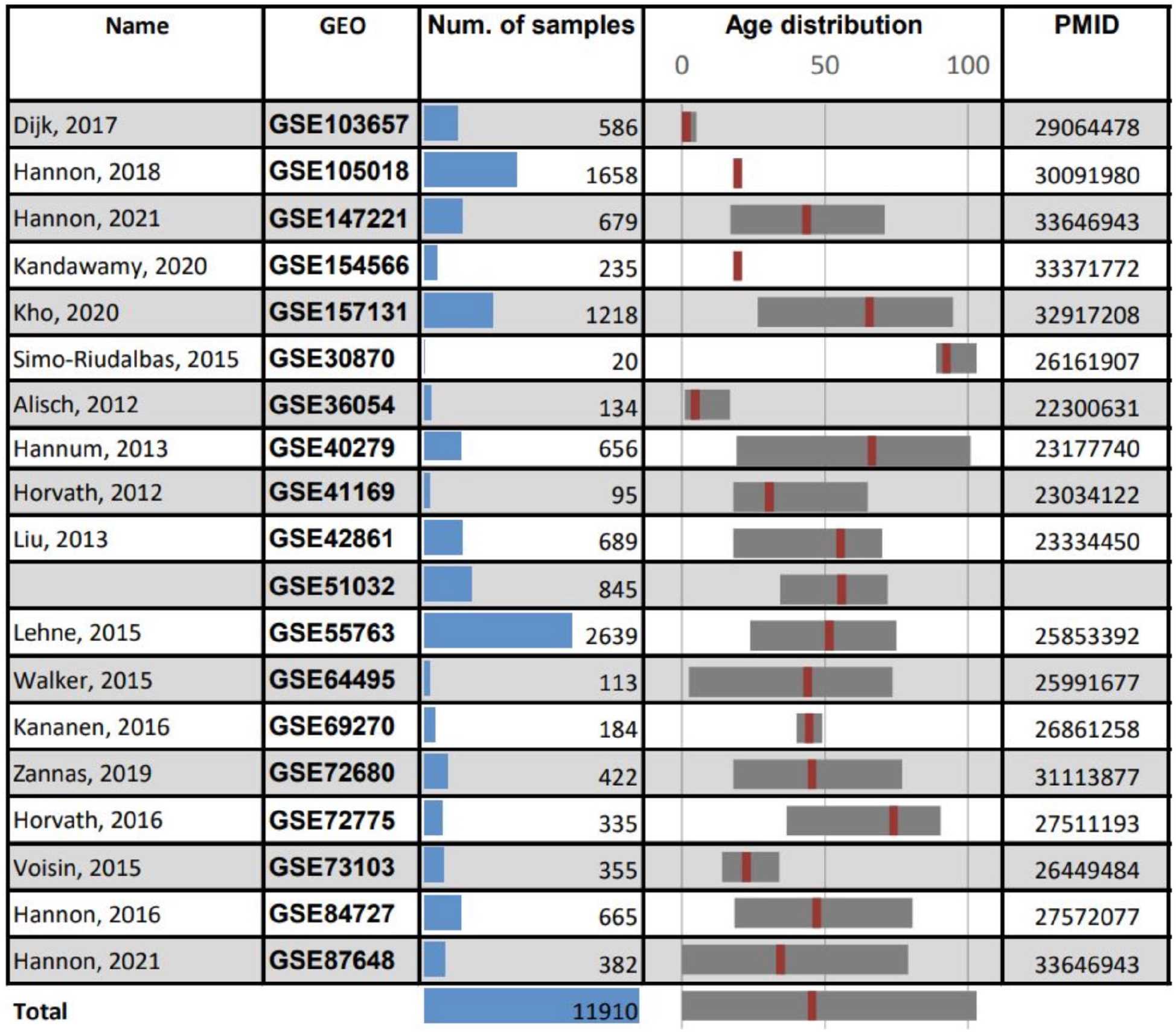
Description of datasets. A total 11,910 blood methylomes were collected from 19 studies. Shown are publication names, GEO accessions, number of samples per dataset, age ranges (median age marked in red), and PMID numbers.

Samples from all other studies were randomly split into a training set cohort (70%, total of n=7,860 samples) and a held-out test set cohort (30%, n=3,385 samples). The train and the test sets show similar age distributions across all datasets. All methylomes were used as published, following their original preprocessing and normalization by various methods^2,21–37^.

### Selection of age-associated CpG sites

We aimed to identify a set of CpGs sites that are highly informative of age. For this, we calculated the correlations between chronological age and methylation levels in train set samples across datasets. Spearman rank correlation was specifically used, to not assume linearity (as in Pearson correlation). Other measures (e.g. Mutual Information) were also tested, but did not reveal additional non-monotonic sites informative of age.

To avoid age-related sites that change at a slow rate, therefore relying on accurate estimations of DNA methylation levels and thus require deep sequencing depth, we preferred CpG sites whose overall methylation gain/loss during adulthood is above 20% (Fig. 2E, Methods). It should be emphasized that samples from the held-out validation set (GSE84727) were strictly excluded from these feature selection processes.

**Figure 2:**
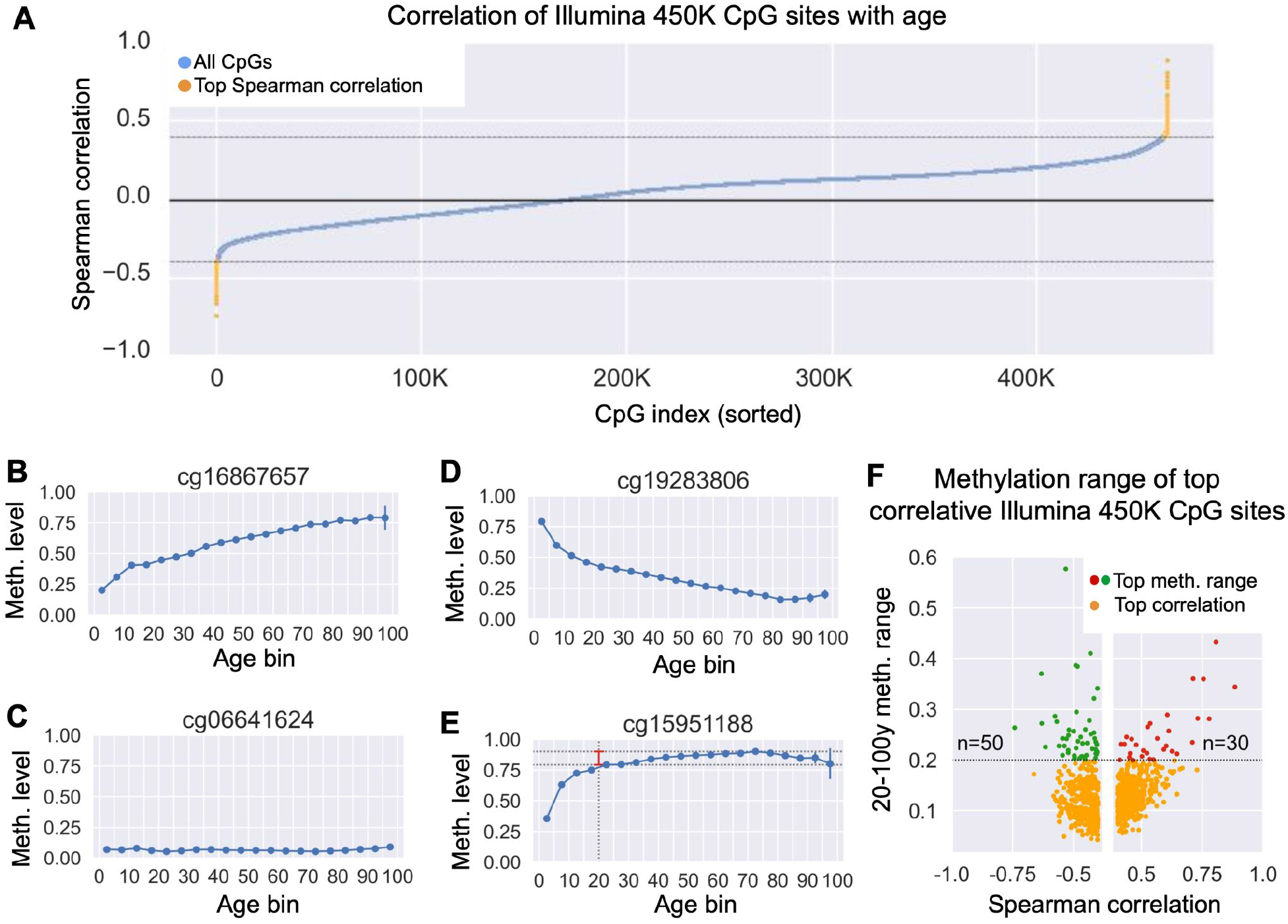
Selection of the age set CpG sites for GP-age. **(A)** Shown are Spearman correlation coefficients for all CpGs from the BeadChip 450K array (blue dots), calculated across training samples ≥20 years old. 964 CpGs with absolute Spearman coefficients ρ≥0.4 were selected for further analysis (yellow dots). **(B-E)** Example of average methylation levels across aging, for four CpG sites. For each 5-year bin, shown are the average methylation levels (blue dots; vertical bars show 95% confidence intervals, CI). Dashed lines and red vertical bar (E) mark the 20-100y methylation range. CpGs with low methylation range (i.e., flat slopes), offer limited age-predictive value and are not included in our age set (e.g. cg06641624 (C), cg15951188 (E)). Conversely, dynamic sites (e.g. cg16867657 (B), cg19283806 (D)) offer high predictive value. **(B)** A CpG with positive age correlation: Spearman ρ=0.88, meth. range=0.34 **(C)** A non-correlative CpG: Spearman ρ=-0.01, meth. range=0.04 **(D)** A CpG with negative correlation: ρ=-0.74, range=0.26 **(E)** Age correlated CpG (ρ=0.4) with meth. range=0.11 (not selected) **(F)** Comparison of correlation (x-axis) vs. the methylation range (y-axis). CpGs with range≥0.2 are shown in red (*ρ*≥0.4, n=30 CpGs) or green (*ρ*≤-0.4, n=50 CpGs). 80 out of 964 correlative CpGs were selected.

Specifically, we analyzed the train set samples across all 485,512 BeadChip 450K array CpGs, retained CpGs with ≤20% of missing values across samples, and calculated their correlation with age (Fig. 2). CpG sites with absolute Spearman correlation ≥0.4 were selected for further analysis (all showing FDR-corrected p-values ≤1e-200). We then computed the methylation range for each CpG, defined as the difference between maximum and minimum methylation values in adulthood, and removed CpGs with span ≤20% (Fig 2E). Overall, our feature selection stages concluded in 80 age-related CpG sites (Fig. 2F). These CpG sites are indicative of age and are robust to sequencing noise (Methods). Selecting alternative thresholds (at each stage of our selection process) did not greatly change the results below.

Aiming to develop a compact model applicable to targeted multiplex bisulfite PCR sequencing, we wished to further reduce the number of CpG sites used. For this, we clustered the candidate CpG sites by their methylation levels across samples to k=30 clusters, and selected the most correlative CpG from each cluster (Methods). Intuitively, CpGs from the same cluster show similar methylation patterns, and therefore contribute little additional information.

### Gaussian Process regression models

We then integrated the k=30 CpG sites into a non-parametric cohort-based Bayesian age predictor, based on Gaussian Process (GP) regression models (Fig. 3A). This is a flexible class of models, estimating the probability distributions over a continuous feature (e.g., chronological age) across multiple (possibly infinite) functions that fit the input data. Unlike parametric regression models (e.g. linear regression models, which assume a constant rate in methylation changes), GPs do not define priors over the parameters of a given set of functions, but rather a distribution over multidimensional functions that are not explicitly defined^38^. Moreover, as these basis functions are not limited to linear functions, they do not require that nonlinear corrections, such as Horvath’s mAge transformation^1^, be applied in the pre-processing step.

**Fig. 3:**
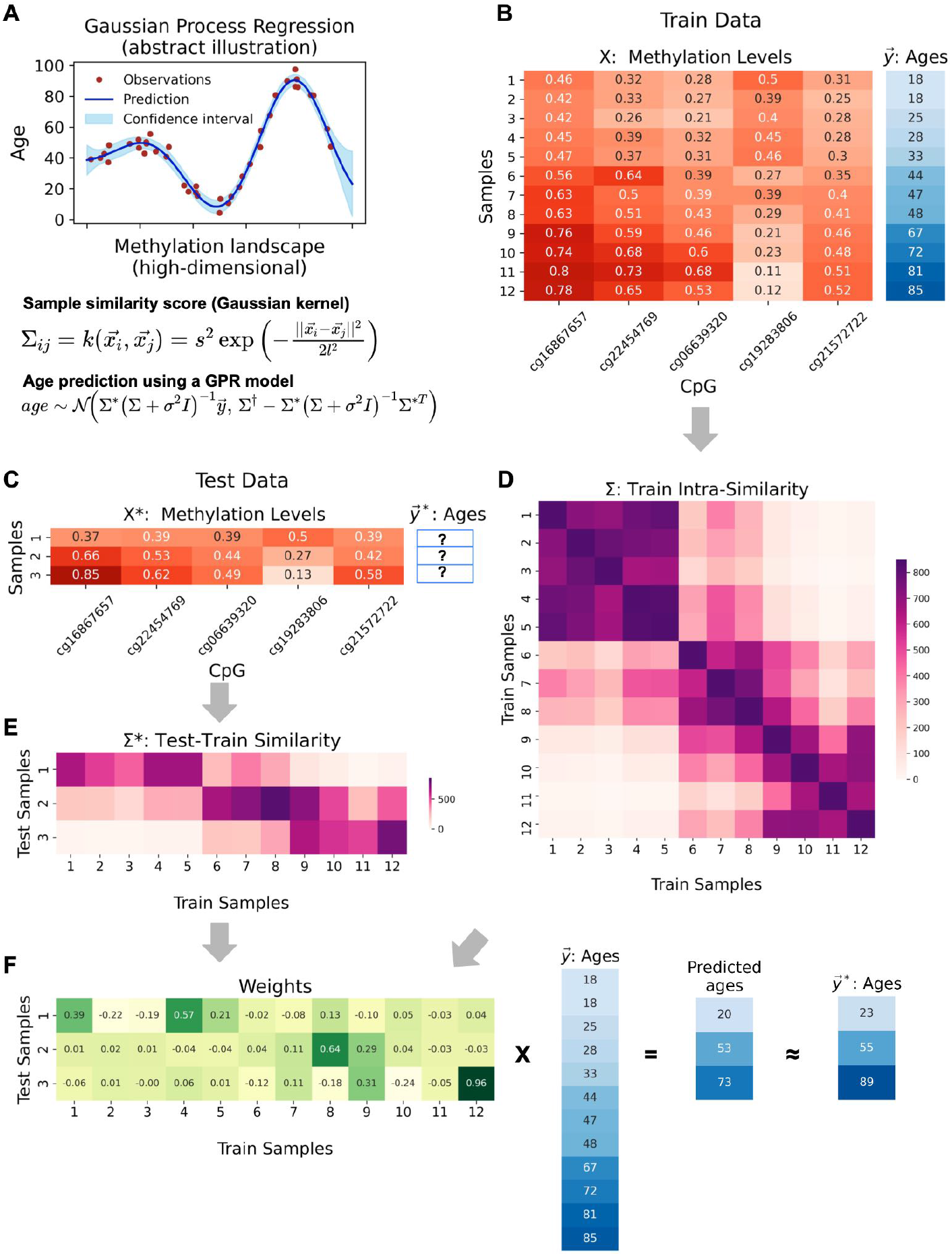
Gaussian process regression model. **(A)** Abstract visualization of a GPR. Shown are example observations (red dots), along with the predictions (blue line) and confidence interval (light blue strip) of the Gaussian process, which is a distribution over possible regression functions from the methylation vectors to chronological age. The Gaussian kernel defining the GP and the distribution of age predictions are shown below. **(B-F): Toy example of age prediction for 3 samples based on 5 CpG sites and a cohort of 12 training samples (from real data). (B)** Shown is a cohort of 12 train samples, containing methylation levels of 5 CpG sites, along with the donor ages. **(C)** Three test samples (methylation vectors over the same 5 CpG sites) are shown. The chronological ages of the samples are unknown. **(D)** The cohort intra-similarity matrix, as calculated with the optimized Gaussian kernel function. **(E)** The similarity matrix between the test and the train set samples as calculated with an optimized Gaussian kernel function. **(F)** The weights assigned to each cohort sample by each test sample are shown, with the resulting final prediction along the real ages of the test samples.

In practice, it is easier to think of GP models in their dual representation: Gaussian kernels are used to measure the similarity across the DNA methylation patterns of train set cohort samples, resulting in the intra-similarity covariance matrix. Given test set samples, the model calculates their similarities to each of the train set samples, which are normalized by the covariance matrix. This results in a weight matrix that associates each test sample to train set samples that show similar methylation patterns. Finally, these samples are weighted and combined to predict the age of each test sample (Fig. 3B-F).

The accuracy of our age prediction model, GP-age, was evaluated by computing the root mean square error (RMSE), which provides a good estimator for the standard deviation of prediction errors. For direct comparison with previous works, we also calculated the median absolute error (MedAE), in years. First, we evaluated the accuracy of the predictions of GP-age with 30 CpGs on the train (7,860 samples from 18 datasets) and held-out test samples (3,385 samples from 18 datasets), resulting in median absolute errors of 2.08 and 2.10 years, respectively (Fig. 4A). RMSE estimations were also very similar between the two sets (3.78 years, train; 3.96 years, test), suggesting that the regression model did not overfit. Importantly, similar accuracy was achieved for the held-out validation set (665 samples from GSE84727), with a median error of 2.24 years, and RMSE of 3.61 years (Fig. 4A). Repeating this procedure with other held-out datasets showed similar results (Supplemental Fig. 2). In agreement with previous works^2,17,39^, the prediction accuracy of GP-age decreases as age increases, as demonstrated by measuring the average prediction error across 5-year bins (Fig. 4B, Supplemental Fig. 3).

**Fig. 4:**
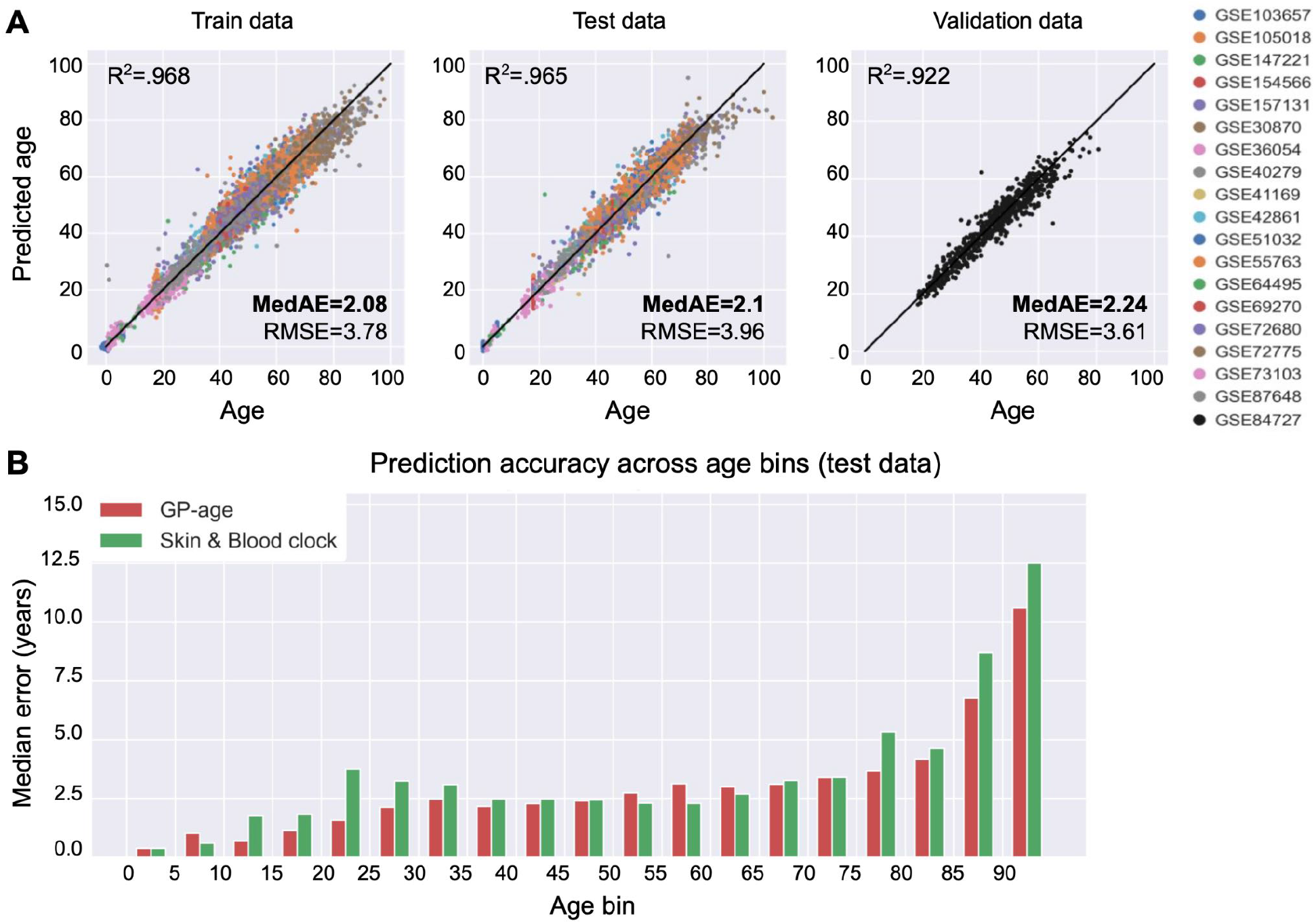
Prediction accuracy of GP-age with 30 CpGs. **(A)** Chronological age vs. predicted age of GP-age with 30 CpGs, across train, test and validation (GSE84727) set samples, yielding a median error of ~2.1 years, and a RMSE of ~4 years, across different datasets (colors). Coefficient of determination R^2^ between prediction and age, RMSE, and MedAE are shown. **(B)** The median error of GP-age with 30 CpGs (red) and the Skin&Blood (green) methylation clocks across different 5-year bins of donor ages.

### Comparison to state-of-the-art age prediction methods

We next turned to analyze chronological age predictions of the test cohorts using published state-of-the-art models, including the 353-CpG multi-tissue clock^1^, the 71-CpG methylation clock by Hannum et al.^2^, and the 391-CpG Skin&Blood clock^10^. As Fig. 5B-C show, these models achieve median errors of 3.9, 4.63, and 2.36 years, respectively, compared to a 2.1-year error by the 30-CpG GP-age model, or a 1.89-year error by the 80-CpG GP-age model. GP-age also outperforms these models on the independent validation set, with a median error of 2.24 years, compared to 6.01, 3.01, and 7.24 years, respectively (Supplemental Fig. 4C). It shall be noted GP-age is more accurate than the Skin&Blood model on young samples (aged 10 through 35) as well as older ones (70 through 95), and is similarly accurate on samples aged 40-45 (Fig. 4B, Supplemental Fig. 3).

**Fig. 5:**
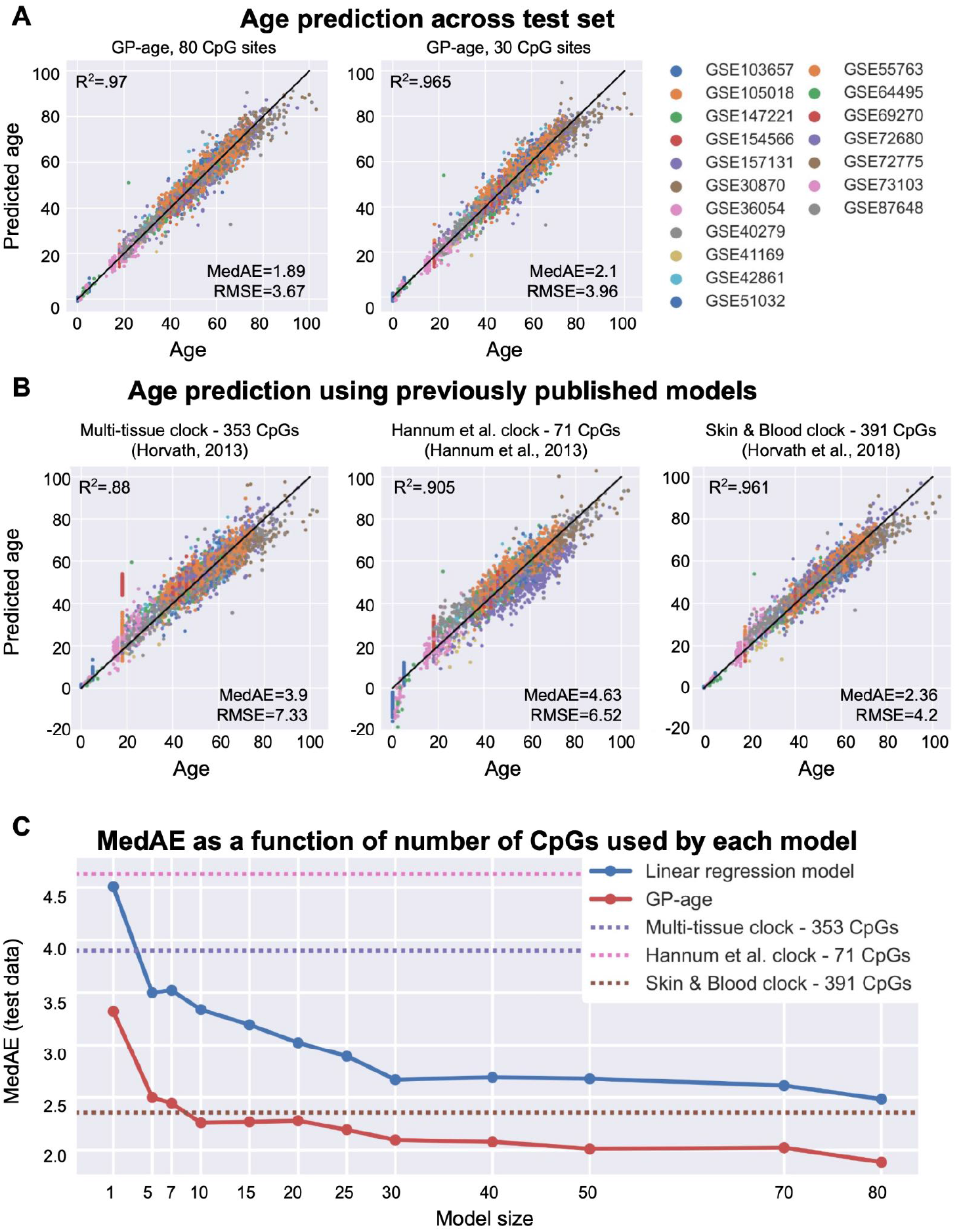
Prediction accuracy of GP-age of different sizes and of state-of-the-art models. **(A)** Age vs. age prediction across test set samples. MedAE of GP-age is ~2 years and RMSE is <4 years, across the different datasets (colors). Increasing the size of the model increases its accuracy. Black line: y=x. **(B)** Age vs. age prediction by previously published models across test set samples. Predictions are less accurate than GP-age, across different datasets. Color legend is the same as in (A). **(C)** Using only 30 CpG sites, GP-age (red) is more accurate than previously published models (horizontal dotted lines), and simpler models trained on the same set of CpG sites (linear regression model in blue). Increasing the model size increases the accuracy.

Other models, including Vidal-Bralo et al.^12^, were also compared to GP-age, and were shown to be less accurate (Supplemental Fig. 6). Unlike most previous models, the clock by Zhang et al.^11^ was trained on an impressively large dataset (~13K samples), including the Lothian Birth Cohorts data^40,41^. While our results show that GP-age offers higher accuracy than all these models, including Zhang et al. (514 CpGs, median abs. error ≥ 8 years) it is possible that this model is tailored to older samples and will be more suitable when applied to special age distributions.

Additional chronological age predictors that use 1,000 CpGs or more, were not compared as they are outside of our scope of this manuscript^11,14,15^.

### Model complexity vs. accuracy

Next, we wished to study how the number of CpGs in the Gaussian Process regression models affects their predictive power. Different model sizes were tested by clustering to k clusters and choosing the most age-correlated CpG from each cluster (Methods). We therefore examined a range of models varying from the full model of k=80 CpG sites, to the single CpG model at k=1. Remarkably, a GPR model with a single CpG (ELOVL2) achieves a median absolute error of 3.3 years, whereas a model with k=10 CpGs outperforms all state-of-the-art models, with a median error of 2.26 years. Overall, the 30-CpG GP-age model allows an optimal tradeoff between prediction accuracy (2.1 years) and compactness, and is comparable to the k=50 and k=70 models (Fig. 5C), as well as the full k=80 CpGs model. Similar results are reported when using the RMSE and MeanAE metrics (Supplemental Fig. 5), and on the held-out validation cohort (Supplemental Fig. 7).

To ensure statistical stability, we applied 10 stratified 4-fold cross-validation runs for each value of k, resulting in an estimated error of <0.01 for the reported median absolute error for each model. In addition to estimating RMSE and median error (50^th^ percentile of predictions), we also estimated the prediction errors at the 25^th^, 75^th^, and 90^th^ percentiles of samples (Supplemental Table 1). Across all percentiles, GP-age consistently outperforms all other models.

### Comparison to non-GPR models and alternations of the feature selection processes

Finally, we compared the GP models of different sizes to linear regression models, similar to the ones used by previously published clocks (Fig. 5C, blue). For all model sizes, linear models were consistently less accurate than the nonlinear cohort-based model, while using the exact same sets of CpGs. To demonstrate the advantage of clustering correlated CpGs, we also compared each model to a GPR model trained on the top-k age-correlated CpG sites (without clustering). For all values of k≤30 CpGs, the clustered sets outperform the top-k sets (Supplemental Fig. 8).

### Retraining GP-age with the validation set

After observing similar prediction accuracy on held-out test data and three independent validation sets (Fig. 4A, Supplemental Fig. 4, Supplemental Fig. 2), we hypothesized that the model is general enough and invariant of different dataset normalizations, and thus combined all 19 datasets for an improved GP-age model. Samples were partitioned into test (30%, n=3,573) and train (70%, n=8,337) sets, and GP-age models trained as described above. Overall, we identified 1,034 age-correlated CpGs (|ρ|>0.4, Supplemental Table 2), 71 of which with methylation range ≥0.2. These were clustered, and updated sets of CpGs were selected (Supplemental Table 3). Intriguingly, for k=30, 27 of the 30 CpGs overlap with the previously selected model, supporting the robustness of the feature selection process. Overall, this model resulted with a median train error of 2.06 years (RMSE=3.74), and a median test error of 2.10 years (RMSE=3.89). Downstream analyses (on held-out test data or external samples), were all performed using these full GP-age models.

### Consistent prediction errors may reflect biological age

To detect consistent prediction errors that may reflect differences between chronological and biological age, we trained three independent GPR models. For this, the 71 age-correlated CpGs were split into three independent groups (composed of 19, 33, and 22 CpGs, from disjoint sets of chromosomes), and GPR models were trained (Supplemental Table 3). The three models show median absolute errors of 2.45, 2.42, and 2.75 years, respectively, on held-out test data. Importantly, the models are as correlated with age as they are with each other (Supplemental Fig. 9).

We next analyzed the pattern of prediction errors across these three independent clocks, and specifically errors consistently larger than the median absolute errors of the models (Fig. 6). Intriguingly, 8.5% of samples were predicted to be younger (“Y”) than their chronological age by all three models, a 6-fold enrichment compared to the expected 1.4% under a null model (binomial p-value ≤ 2e-133). Similarly, 7.5% of samples are consistently predicted as older (“O”), a 4.5-fold enrichment compared to the 1.7% expected (p-value ≤ 2e-88). Enrichment was also observed for the consistently age-matching (“M”) predictions (21% observed, 12.5% expected, p-value ≤ 3e-46). These results suggest that only a small fraction of consistently biased samples are random, while the remaining, 83% of consistently younger predictions, and 77% of consistently older predictions, reflect true deviations between chronological and biological ages.

**Fig. 6:**
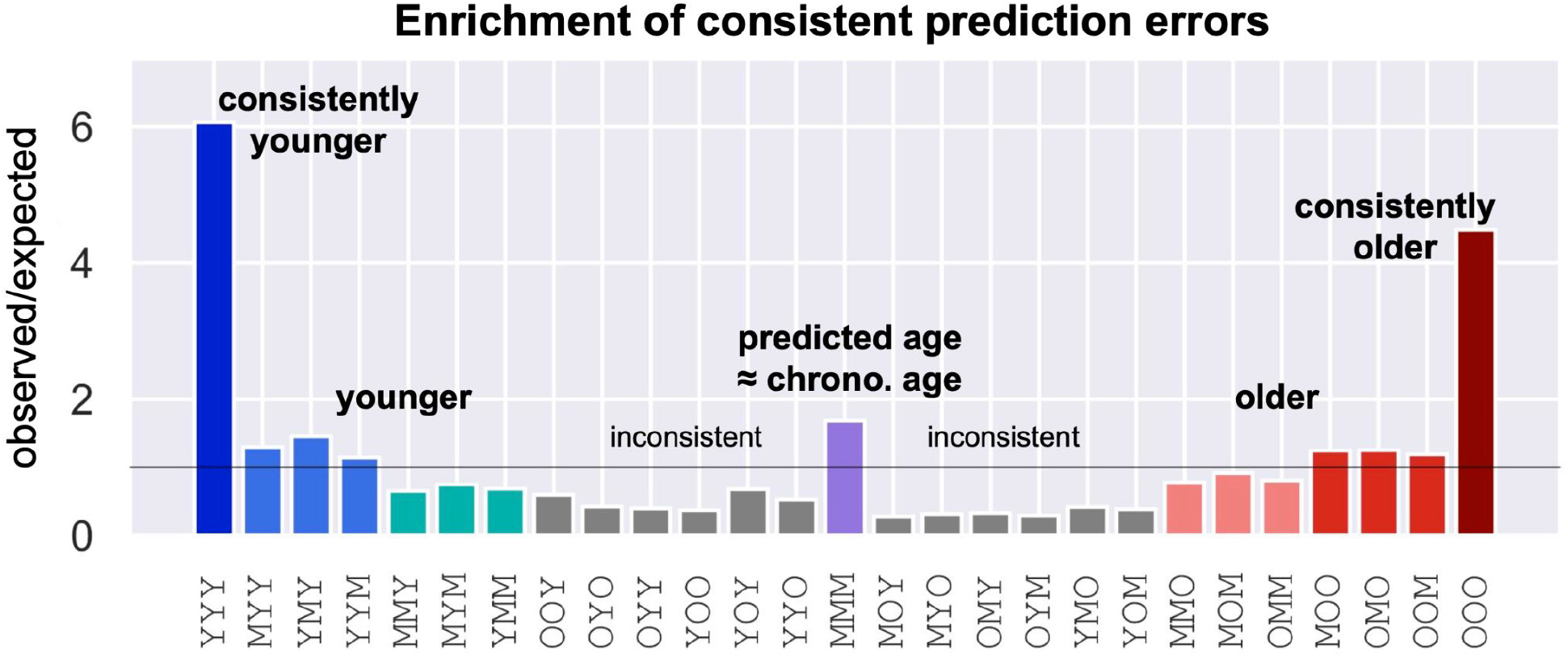
Consistent prediction errors across independent GPR models suggest biological variance. Enrichment of prediction error patterns across three independent clocks. Y: predicted as younger than real age. O: predicted as older. M: prediction matches age, within the median abs. error. 8.5% of samples are consistently predicted to be younger than real age (blue, 1.4% expected, fold change of 6.06), 7.5% of samples are consistently predicted as older (burgundy, 1.7% expected, FC=4.48), and 21% are predicted within median abs. error by all three models (lilac, 12.5% expected, FC=1.68).

### Methylation trends of clock CpG sites

To better understand the age-related dynamics of the 71 CpG sites in GP-age, we divided the CpG sites to those gaining methylation over age (n=33), and those losing methylation (n=38). Intriguingly, while both positively and negatively age-correlated sites were identified, we observed a small enrichment for CpG sites that gain methylation with age (Fig. 7A). Accordingly, when training two separate models, the epigenetic clock that uses positively correlated (“gaining”) sites shows a higher accuracy than the one based on negatively correlated ones (“losing”) (Fig. 7B, Supplemental Fig. 10), with median errors of 2.54 and 3.45 years, respectively. Interestingly, the gap between the two clocks becomes smaller when the ELOVL2 CpG site cg16867657, is excluded from the positive set, leading to a median error of 2.83 years. These results suggest that age-related changes in DNA methylation involve both hyper- and hypo-methylation.

**Fig. 7:**
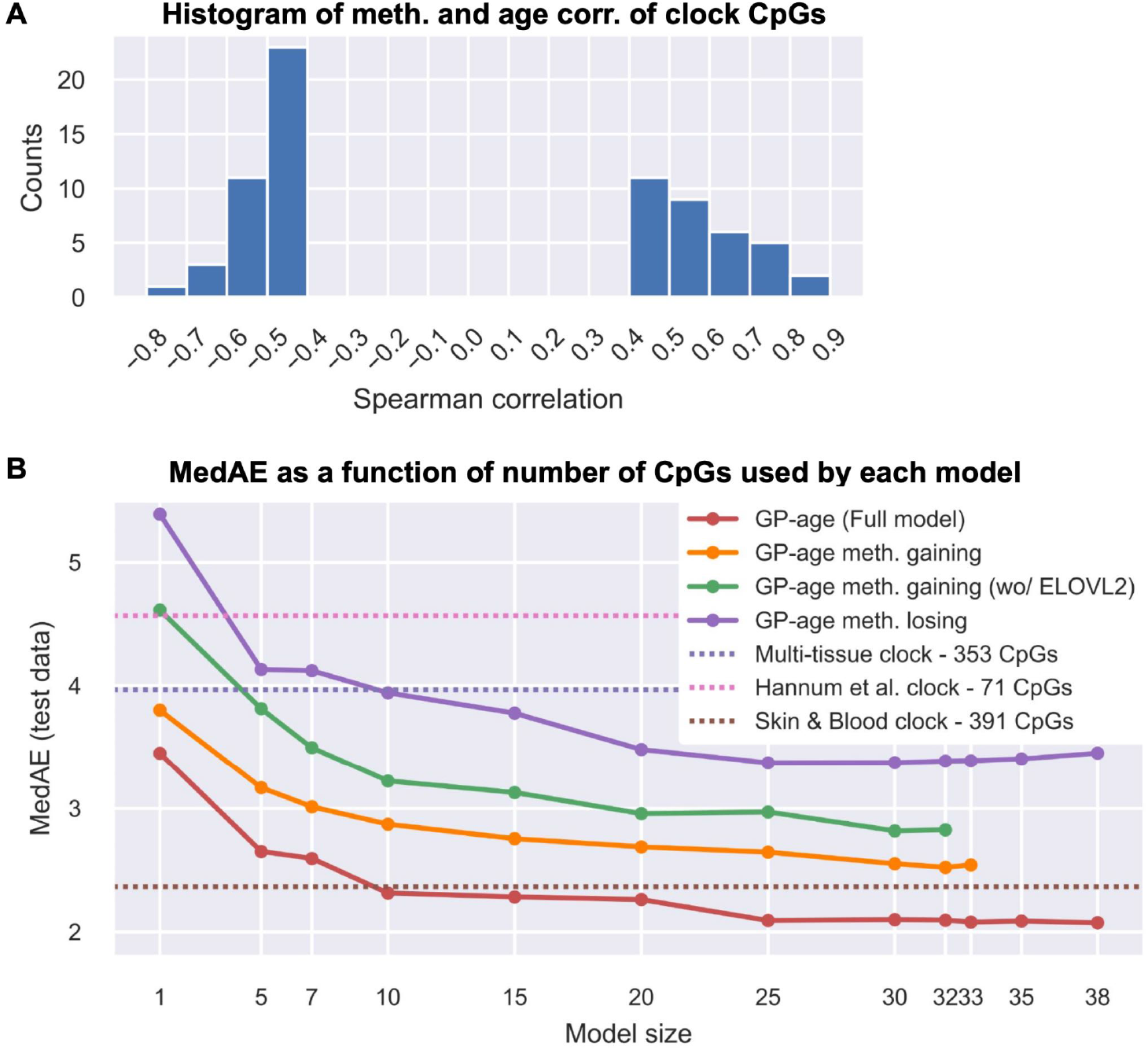
Methylation-gaining CpGs are more correlated with age than methylation-losing CpGs. **(A)** Histogram of the Spearman correlation of methylation and age of the GP-age clock CpG sites. **(B)** Model trained with only methylation-gaining CpG sites (orange) is more accurate than models trained with methylation-losing CpG sites (purple). This is still observed when the ELOVL2 CpG is excluded (orange). The full GP-age model outperforms these models (red).

### Accurate age predictions from whole-genome bisulfite sequencing data

Encouraged by these results, we wished to establish our 30-CpG epigenetic clock in sequenced-based DNA methylation data from blood, in addition to methylation array data as previously shown. Unlike methylation arrays, whole-genome bisulfite sequencing (WGBS) methylation data is relatively shallow with typically no more than 30 sequenced reads (fragments) covering each CpG (30x). Yet, neighboring CpG sites are also sequenced and could be an additional source of data, at least in genomic regions with block-like methylation patterns^42–45^. We therefore turned to test the performance of GP-age on such data.

GP-age with 30 CpG sites was applied to two blood WGBS datasets. Initially, average methylation was calculated at k=30 CpGs of the GP-age model, and age was directly predicted. As Fig. 8 shows, this resulted in a median error of 3.0 years (RMSE=6.10) on buffy coat methylomes sequenced by Jensen et al. at 29x depth from n=7 donors aged 20-4746. We also applied GP-age to a set of n=23 deeply sequenced (83x) leukocyte methylomes of donors aged 21-75, recently published by us^45^, resulting in a median error of 3.55 years (RMSE=4.92).

**Fig. 8:**
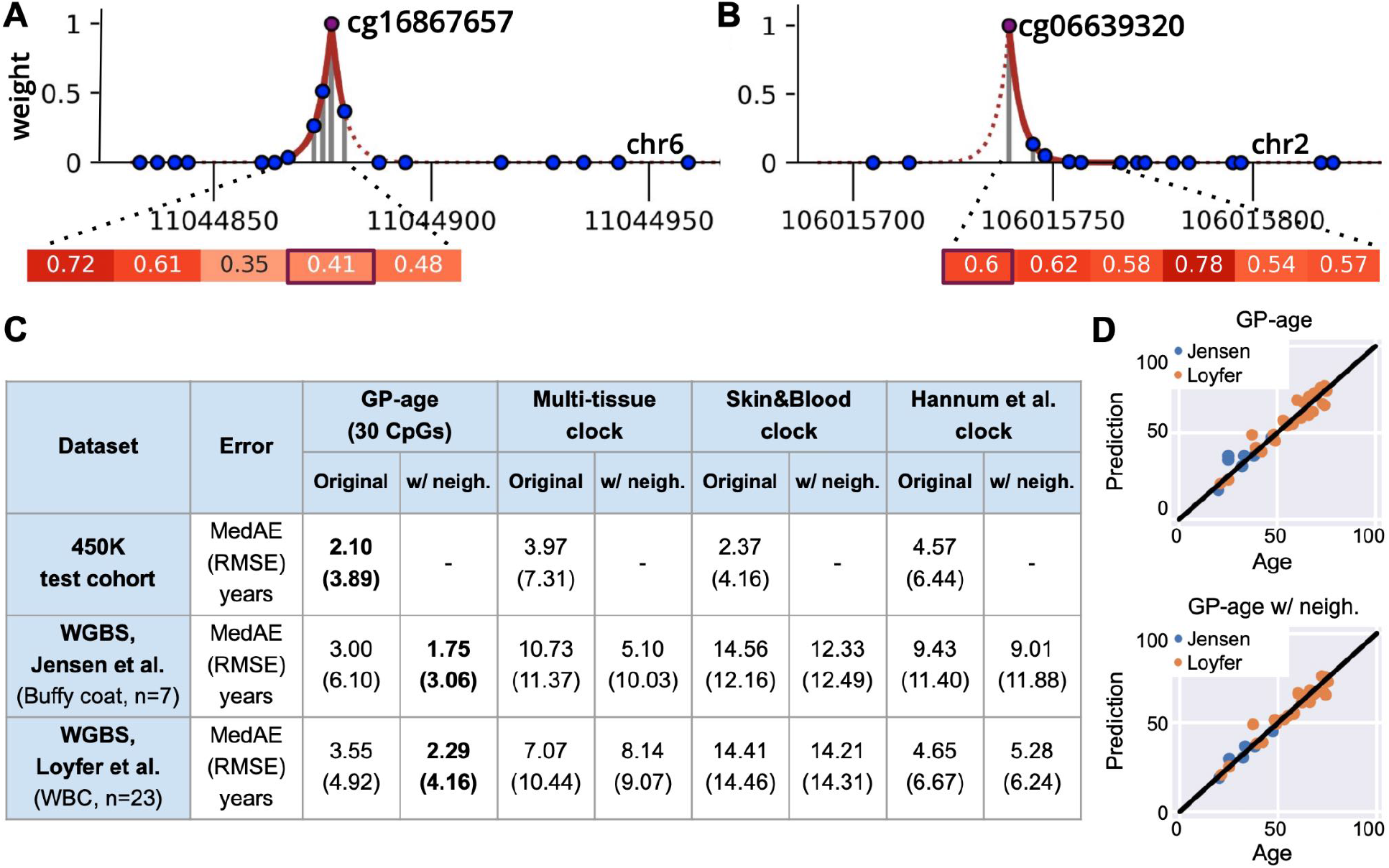
Weighted average of neighboring CpG sites in WGBS data. **(A-B)** Genomic region of a CpG was defined with a genomic segmentation into methylation blocks. A Laplace kernel (dotted red line) was used to assign weights to the CpGs in the neighborhood (solid red line). Methylation levels of CpG sites in the neighborhood are shown. **(C)** RMSE and MedAE errors of GP-age and reference models (with and without neighbors), across datasets. The lowest error for each dataset is marked in bold. **(D)** Age prediction across the two sequencing datasets. Top: Using a single-CpG resolution methylation level for each CpG from the age set. Bottom: Using a Laplace kernel for a methylation level estimation by a weighted average of methylation levels of neighboring CpGs.

Encouraged by these results, we wished to incorporate the methylation values of neighboring CpG sites, to compensate for the relatively low coverage of the data. For this, we segmented the human genome (hg19) into homogenous methylation blocks^45^, and averaged the target CpG with surrounding sites, weighted using an exponentially decaying Laplace kernel (Methods). As Fig. 8 shows, this further improved GP-age’s predictions on WGBS datasets, yielding a median error of 1.75 years on the Jensen et al. dataset^46^, and 2.29 years on the Loyfer et al. dataset^45^. The difference in accuracy between the two datasets could be explained by the different ages in the two datasets (Supplemental Fig. 11B-D). This again is consistent with the previously reported decrease in the accuracy of epigenetic clocks as age increases.

Notably, previously published array-based methylation age models^1,2,10^, all showed higher errors of 5-15 years for the two datasets (Fig. 8C). We reason that this is partly due to age-correlated CpG sites that do not change greatly overall (low methylation range), and are therefore hard to approximate at WGBS sequencing depths. Other age prediction models used pyrosequencing data^13,17,47^ at few CpG sites, mostly for forensic use. GP-age with 30 CpGs outperforms these models as well (Supplemental Fig. 11A).

## Discussion

In this article we present GP-age, a non-parametric cohort-based chronological age prediction model, and compare it to previously published state-of-the-art models. While other epigenetic clocks were developed for different tissues, or as multi-tissue predictors, in this work we focus on whole blood, as it is easily accessible. Future work may be to apply a similar method to develop Gaussian Process-based epigenetic clocks for other tissues.

GP-age uses a cohort of 11,910 blood methylomes, measured using Illumina BeadChip 450K/EPIC methylation arrays. These are made available as a resource to the methylation age community. Samples from various cohorts were merged, and split into test and train sets. Sets of non-redundant age-correlated CpGs were then selected. In this cross-cohort analysis, we specifically did not renormalize samples from different datasets, so CpG sites with batch differences were implicitly selected against. We then trained a non-parametric Gaussian Process regression model, which uses these CpGs to compare a query sample against the train set cohort, find similar methylomes, and predict the query age based on train set ages and intra-cohort dependencies.

As we show, A 30-CpG GP-age model achieves a median absolute error of 2.1 years across 3,573 held-out test samples, outperforming state-of-the-art methods (on the same data). An even more compact model, consisting of only 10 CpGs, is comparable to state-of-the-art clocks with a median error of 2.26. Similar results were achieved on parallel GP-age models, for which one of the datasets was considered as a validation set, and its samples were excluded from feature selection and model training (Fig. 4A and Supplemental Fig. 2, showing three different such validation sets).

Depending on the desired model size and accuracy, 10-, 30- or 71-CpG GP-age models are suggested for age prediction, as these models provide a good tradeoff between compactness and accuracy. As we further show, the model is also applicable to next-generation sequencing data, where a Laplace kernel is used to augment the methylation levels of the age prediction CpGs by their neighboring CpGs. This resulted in a similar prediction accuracy of ~2 years.

It shall be noted that previous studies presented highly compact chronological epigenetic clocks, sometimes involving as few as three CpGs. Nonetheless, these compact models presented inferior accuracy, with median absolute errors of 5-21 years on our Illumina 450K and sequencing data^12,13,16–18,47^. Conversely, we provide an epigenetic clock that is more accurate than commonly used models^1,2,10^, and at the same time compact enough to allow direct measurement using multiplex targeted PCR, making these models simpler and more accessible compared to DNA methylation arrays.

Overall, GP-age predictions of chronological age outperform current state-of-the-art models while using fewer CpG sites, thus opening the way for various applications in aging, forensics, transplantations or more, using low-cost targeted PCR sequencing data. These results were achieved by three independent means. First, we selected a set of CpG sites whose average methylation changes with age. As we showed, using these sites to train a linear regression model, similar to the ones used by Horvath^1,10^ and Hannum et al.^2^, already achieves a median absolute error of 2.70 years. The compactness of the model is achieved by selecting one representative from each cluster, thus minimizing similarities between model CpGs. Second, we assembled a large training cohort which allows cohort-based models to identify similar methylomes for each query set. Third, Gaussian Process regression models add accuracy by not being limited to a fixed number of neighbors (as in KNN), and use intra-cohort similarities to further determine how these samples are weighed. Thus, through a more complex assignment of weights to train samples, GP-age utilizes more information from the cohort, resulting in higher accuracy.

Several of the CpG sites that were automatically selected by our model are known to be associated with age-related genes or have been previously included in epigenetic clocks. Most notable is ELOVL2 ^2,48,49^, single-handedly providing a median absolute error of 3.3 years in our cohort-based algorithm. Intriguingly, exclusion of ELOVL2 resulted in a median error of 2.49 years using a GP-age model with 10 sites (and 2.28 years using 30 sites). Additional genes were previously associated with aging, including FHL2^2,10,48,49^, OTUD7A^2,49^, CCDC102B^2,49^, TRIM59^2,10,50^, RASSF5^51^, GRM2^52^, ZEB2^53,54^, Zyg11A^49^, TP73^55^, IGSF11^56^, MARCH11^57^, SORBS1^52^, ANKRD11^58^, and EDARADD^1,2,10,59^. The remaining CpGs, including cg20816447 (CC2D2A), cg06155229 (PMPCB), cg06619077 (PDZK1IP1), cg19991948 (TIAL1), cg22078805 (FAM171A2), and cg17621438 (RNF180), were not, to the best of our knowledge, previously associated with aging and should be further studied. The set of CpG sites used by GP-age consists of both methylation-gaining and methylation-losing sites. Intriguingly, a GPR model that uses only methylation-gaining CpGs predicts age better than a GPR model that uses only methylation-losing CpGs (Fig. 7). This observation is still valid when the ELOVL2 CpG is excluded, and raises questions regarding the biochemical processes that underlie the changes of the epigenetic landscape with age.

As previously reported^1,10,12^, changes in age-correlated CpGs reflect cellular changes upon aging, and cannot be explained by quantitative changes of different cell types in the blood (Methods, Supplemental Fig. 12). Furthermore, differences between predicted and chronological age could reflect a biological signal rather than predictions inaccuracy, as we demonstrated with coordinated prediction errors from three independent models (Fig. 6). Thus, although GP-age was not trained as a biological age predictor per se^19,20^, the high availability of DNA methylation data (e.g. the large cohort presented here), opens opportunities to directly study the molecular mechanisms of aging, as well as variation across individuals, which could not be conveniently addressed using current biological age clocks or datasets that involve additional biochemical features.

Age prediction errors across multiple samples are often summarized as the median absolute error (MedAE), which could be somewhat different than the root mean squared error (RMSE). While the median error provides an upper bound of the error for half the samples (regardless of the other half), the RMSE score provides the standard deviation of predictions across all samples. The differences between RMSE and MedAE scores are observed in the prediction statistics of both the array-based models (Fig. 8C) and the targeted PCR-based models (Supplemental Fig. 11A). For GP-age and the array-based models, these differences are partially explained by the lower accuracy achieved for older ages (Fig. 4B), in agreement with previous studies^2,17,39^. Importantly, GP-age outperformed state-of-the-art models in both measures.

We show that both GP-age and the Skin&Blood clocks are more accurate on younger samples, with the highest improvement by GP-age achieved on ages 10-45 and 70-95 (Fig. 4B). Importantly, specific models explicitly trained on target age groups did not improve the prediction accuracy. Similarly, female- or male-only clocks have not obtained higher accuracy on held-out test data (data not shown). These results suggest that the methylation clock presented here reflects universal processes involved in aging.

GP-age also showed increased accuracy when predicting age from next-generation sequencing data. Here we augment this information by incorporating neighboring CpGs, with their importance decaying exponentially the further they are from the target CpG site (Fig. 7). This, combined with the small set of CpGs, suggests that GP-age could be used to predict chronological age from blood samples using sequencing data, including genomic DNA enriched by hybrid-capture panels for specific age-related regions, or even multiplex targeted-PCR data, which are more accessible than methylation arrays, and with shorter turnaround time.

As we show, methylation arrays provide extensive information regarding the methylation landscape of a given sample, but for the purposes of chronological age estimation, a small set of CpGs suffices. Future research should validate GP-age on targeted PCR data. Notably, GP-age shows higher accuracy than forensic age prediction models^13,17,47^ when tested on identical sequencing data.

Notably, the improved predictions in WGBS samples using information from neighboring CpG sites, suggests that aging-related epigenetic changes occur - at least for some genomic loci - at DNA methylation blocks^42,45^, rather than at isolated independent CpGs (as may seem from DNA methylation array data). Further, this raises questions regarding the underlying processes of epigenetic aging. A possible research direction may be the examination of the recruitment of methylases and demethylases to specific loci, their dynamics, their processivity across neighboring sites, and how these change with age. Most importantly, we show that NGS data from 30 CpG sites at a sequencing depth of 30x achieves <2 year accuracy. This implies that ~1,000 DNA molecules at regions carefully selected, are enough for accurate age prediction.

The use of GP-age could involve a variety of applications, including forensic profiling, transplantation medicine, and health monitoring. While the models presented here were trained and tested on blood-derived methylomes from healthy humans, future works could further expand this approach to other species, tissues, or clinical conditions. Most importantly, its unique simplicity and shorter turnaround times could facilitate longitudinal studies that will shed light on the molecular processes underpinning human aging.

## Methods

### Data and code availability

The data analyzed in this study were deposited at Gene Expression Omnibus (GEO) under accession GSE207605. A standalone implementation for age prediction from array methylomes is available at https://github.com/mirivar/GP-age or from the lead contact.

### 450K Dataset

We assembled a large whole-blood DNA methylation dataset by combining 11,910 blood-derived methylomes from 19 publicly availables individual datasets, measured on the Illumina 450K or EPIC array platform (Fig. 1). All donors included in our dataset were healthy, with ages spanning the range of 0-103 years with a median of 43 years. The partition into the train and test sets was performed randomly, with 30% of the assembled dataset (3,573 samples, median age of 43 years) assigned for test and held out, and the remaining 70% (8,337 samples, median age of 44 years) were used for feature selection and model training. For the initial analysis, one dataset (GSE84727) was held-out and used for validation. Normalized beta values from the original datasets were downloaded from GEO and used without additional normalization or batch correction, to facilitate use of the GP-age model by future datasets. Missing values of each CpG site were imputed with the average beta value of that CpG across other samples, using SimpleImputer from the python package *scikit-learn^60^* (version 0.24.2).

### CpG feature selection

Low-quality CpG sites, with >20% of missing values across train samples, were removed. The Spearman correlation between methylation levels and age was calculated independently for each CpG site from the Illumina 450K platform, across train samples. For a more robust correlation estimation, in order to avoid outlier effect by young (<20 years) samples, datasets including exclusively young donors (GSE154566, GSE105018, GSE36054 and GSE103657) were excluded from correlation analysis (Supplemental Fig. 1). Overall, 964 CpG sites showed an absolute spearman ρ≥0.4 (or 1,034 sites across all 19 datasets, including GSE84727), and were retained for downstream analysis.

Next, the range of methylation levels was calculated for each CpG independently by calculating the methylation average in adults (≥20) in 5-year bins, and calculating the difference between the maximal and minimal values. CpGs sites with range <0.2 were excluded, yielding a set of 71 candidates (when the validation set GSE84727 was included; 80 when not). These were clustered using the spectral clustering algorithm by Ng, Jordan and Weiss^61^, with ke{1,5,7,10,15,20,25,30,40,50,70,80} clusters, and the top correlated CpG was selected from each cluster. For most analyses, we used k=30, but k=10 and k=80 are also reported.

### Gaussian Process regression

A Gaussian Process regression (GPR) model was developed to predict the age of donors given their methylome. GPR is a flexible non-parametric Bayesian approach for regression. In our model, the inputs are blood-derived methylomes over k=30 CpG sites, and the outputs are the ages of the donors. The model was trained using the python *GPy* package (version 1.10.0)62, with the default hyper-parameters adjustments.

A Gaussian Process (GP) is a probability distribution over possible functions that fit a set of points. Formally, it is a collection of random variables, any finite number of which have a joint Gaussian distribution^38^. Given a finite set of input points:

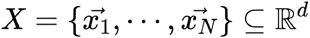

a mean function:

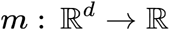

and a covariance function:

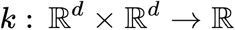

A GP *f* can be written as:

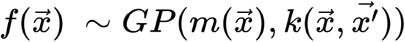

if the outputs 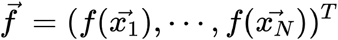 have a Gaussian distribution described by: 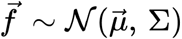, where 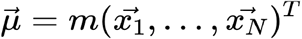 and 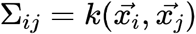. The mean function is usually assumed to be the zero function, and the covariance function is a kernel function chosen based on assumptions about the function to be modeled. In our modeling, we used the commonly used RBF kernel function, defined as:

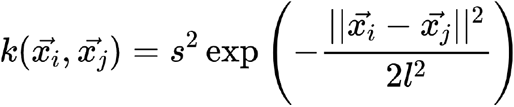

where *s*^2^ is the variance hyper-parameter, and *l* is the length-scale hyper-parameter which controls the smoothness of the modeled function, or how fast it can vary.

In summary, with noise-free observations, the training data comprises of input-output pairs such as:

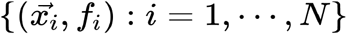

where the inputs are 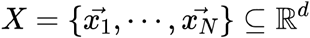 and the outputs are distributed according to a normal distribution: 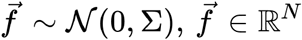

Often, the output variables are assumed to further include some additive Gaussian noise *η*. In which cases the training data can be written as:

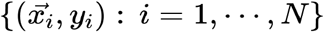

whereas 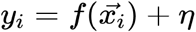, with 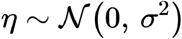. Under these assumptions, where the noise is independent and of equal variances, and outputs could be written as 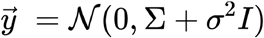. To fit a GP for a regression task, the hyper-parameters of the model 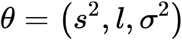 are optimized with respect to the training data. If the mean of the GP is set to zero, Python’s *GPy* package estimates the hyper-parameters by minimizing their negative log marginal likelihood:

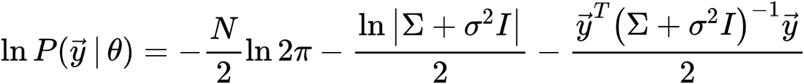

Given a test point 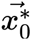, its output distribution is defined by 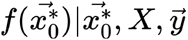, and can be analytically derived. From the definition of a Gaussian process, the finite set 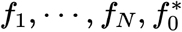 are jointly distributed as:

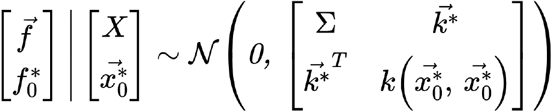

where

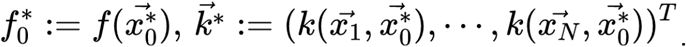

Adding the Gaussian noise to the observations, the finite set *y*_1_,…,*y_N_,f*\* are jointly distributed as:

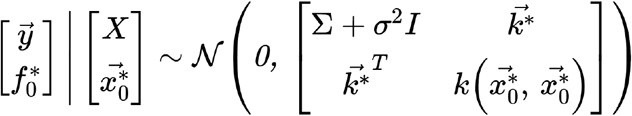

Conditioning the joint Gaussian prior distribution on the observations gives the following conditional distribution:

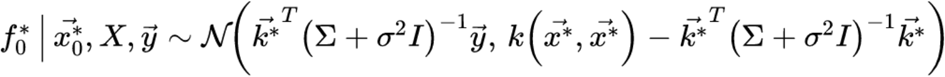

For a set of new samples, instead of a single test sample:

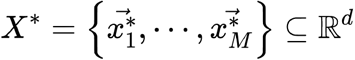

a prediction can be made by taking the mean of the well-defined conditional distribution:

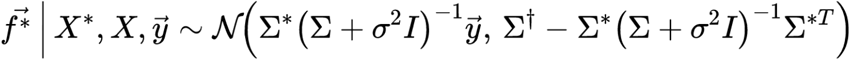

where

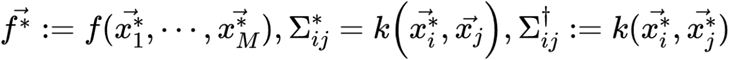

Thus, given the training data, the distribution of predictions of a new point or set of points is given by a closed analytical form. In our model, the inputs are methylation vectors, and the outputs are the donor ages. The mean of the distribution can be used as the final prediction of the regression model.

It shall be noted that the mean term of the conditional distribution of the new output variable derived from the joint Gaussian distribution could be viewed as a weighted sum of the train set ages (Fig. 3). Here, the weights are based on the covariance between the input sample and the train data samples, then multiplied by the inverse of the covariance of the train set cohort data (with Gaussian noise added). Intuitively, this procedure gives higher weights to train set samples with methylomes similar to the query, but penalizes train set samples that are similar to each other, as they do not provide additional information. That way, the GP builds a nonlinear relationship between input vectors and output variables.

### Comparison to previously published 450K-based models

Previously published chronological age predictors^1,2,10^ were tested on our test set using the R *methylclock* package (v 0.7.7)63. An intercept of −5.5 was added to the Hannum et al. clock, for calibration. The Zhang et al. clock^11^ was tested with their provided code, and the Vidal-Bralo et al. clock^12^ was tested with a linear regression model using their published coefficients.

### Training of other regression models

Using the same CpG sites as of GP-age, we trained linear regression models and KNN models for chronological age predictions. The models were trained with LinearRegression (default parameters) and KNeighborsRegressor (n_neighbors=3, p=1, weights=’uniform’) from the python package *scikit-learn* (version 0.24.2)^60^, accordingly.

### Stratified 4-fold cross validation

To check the robustness of GP-age, we performed 10 repetitions of stratified 4-fold leave-one-out cross validation. The samples were divided by binning the donor ages into 5-year bins, and in each repetition, each bin was divided into four subgroups. A single subgroup was retained for validation each time, resulting in four different models for each repetition. The errors across repetitions of cross validations were logged. The mean error and its 95% confidence interval were calculated, using a t-distribution with n-1 degrees of freedom.

### Independent models and coordination of prediction errors

The 71 age-correlated were divided into three groups, such that the CpGs in each group are from distinct chromosomes. Three GP-age models were then learned, from the training samples, as described above. The median absolute error was then determined for each clock, as well as percent of training samples with a greater positive error (over-estimation, ‘O’, average of ~26% of training samples across three clocks), or a greater negative error (under-estimation, ‘Y’, average of ~24% across three clocks). These percentages were used to estimate the expected frequency of each pattern of prediction errors across three clocks. Binomial distribution was used to estimate the statistical significance of enrichment at specific patterns (OOO, YYY).

### Blood deconvolution

All 11,910 methylomes were deconvoluted using our previously published human DNA methylation atlas^44^ (https://github.com/nloyfer/meth_atlas), including seven blood cell types. Proportions of each cell type were then grouped across samples in 5-year bins.

### WGBS data for validation

Two whole-genome bisulfite sequencing datasets have been used in our study. First, we used a dataset published by Jensen et al.^46^ in their study of cell-free DNA in pregnant women. The dataset contains, along with other samples, the methylation levels of 7 samples isolated from maternal buffy coat cells. Bam files were downloaded from dbGaP (accession number phs000846), and analyzed by *wgbstools* using the bam_to_pat function^64^. Second, we analyzed a dataset published by us^45^, including 23 WGBS white blood cells (WBC) samples from healthy donors.

### Testing on WGBS

Single CpG resolution of WGBS dataset was obtained with *wgbstools*, a computational suite we recently developed^64^, using the beta_to_450K function which calculates the average methylation levels of each CpG. These were then analyzed by GP-age for age prediction. We also augmented the methylation level estimation at each target CpG *x* by considering neighboring CpGs *x_i_* within the same DNA methylation block^45^, and averaging their methylation levels using an exponentially decaying Laplace kernel:

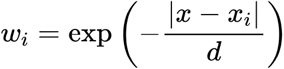

with *d* being a parameter controlling the length scale of effect of CpG neighbors. Here, we used *d*=3.

## Acknowledgements

We thank Nir Friedman, Netanel Loyfer, Alon Appleboim, Josh Moss, Mor Nitzan, Yair Weiss, Roy Friedman, Michael Hassid, and members of the Kaplan and Dor labs for helpful discussions and comments.

## Authors’ contributions

M.V., B.G., Y.D., R.S., and T.K. conceived and designed this research. G.H. and M.V. compiled all data. M.V. and G.H analyzed the data. M.V. and T.K wrote the paper.

## Supplementary Figures

**Supplemental Fig. 1:**
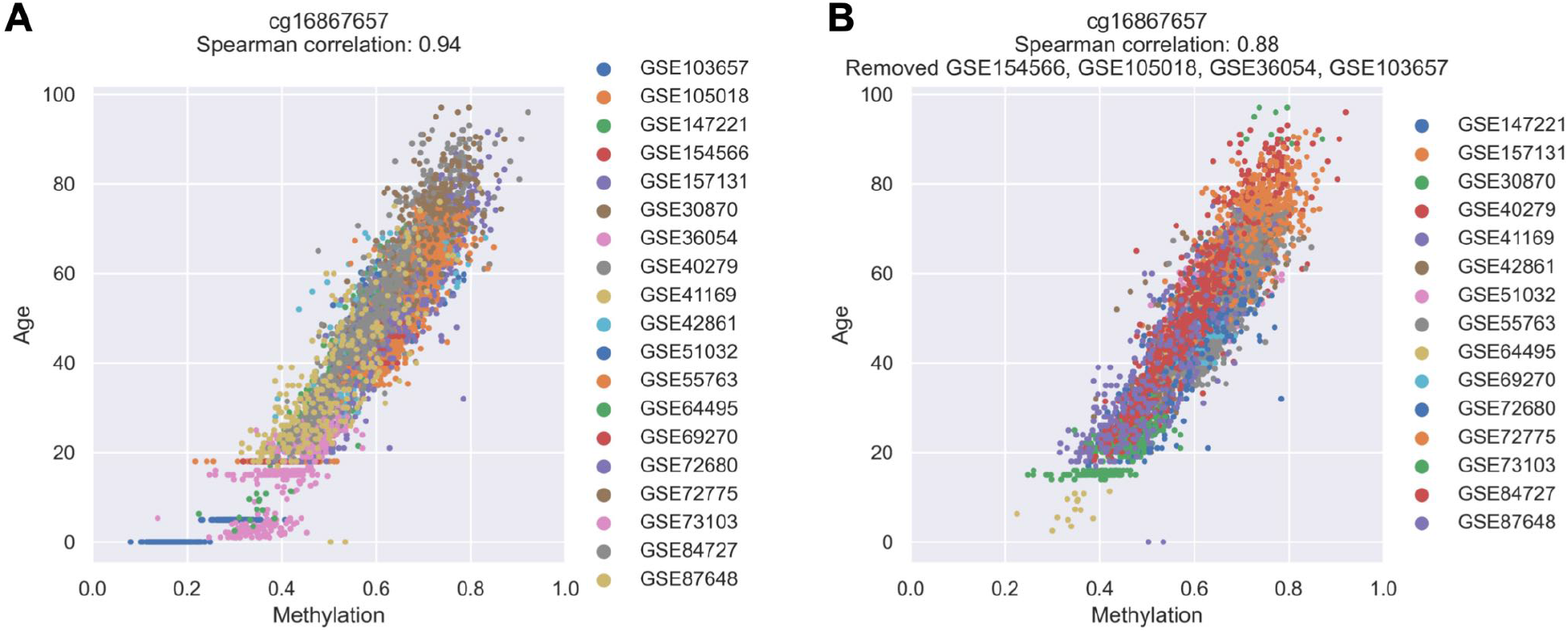
Removal of datasets exclusive for young (<20) donors for correlation analysis. **(A-B)** Shown are average methylation levels of cg16867657 in different samples across all datasets (A); or after removal of the young age datasets: GSE154566, GSE105018, GSE36054, and GSE103657 (B).

**Supplemental Fig. 2:**
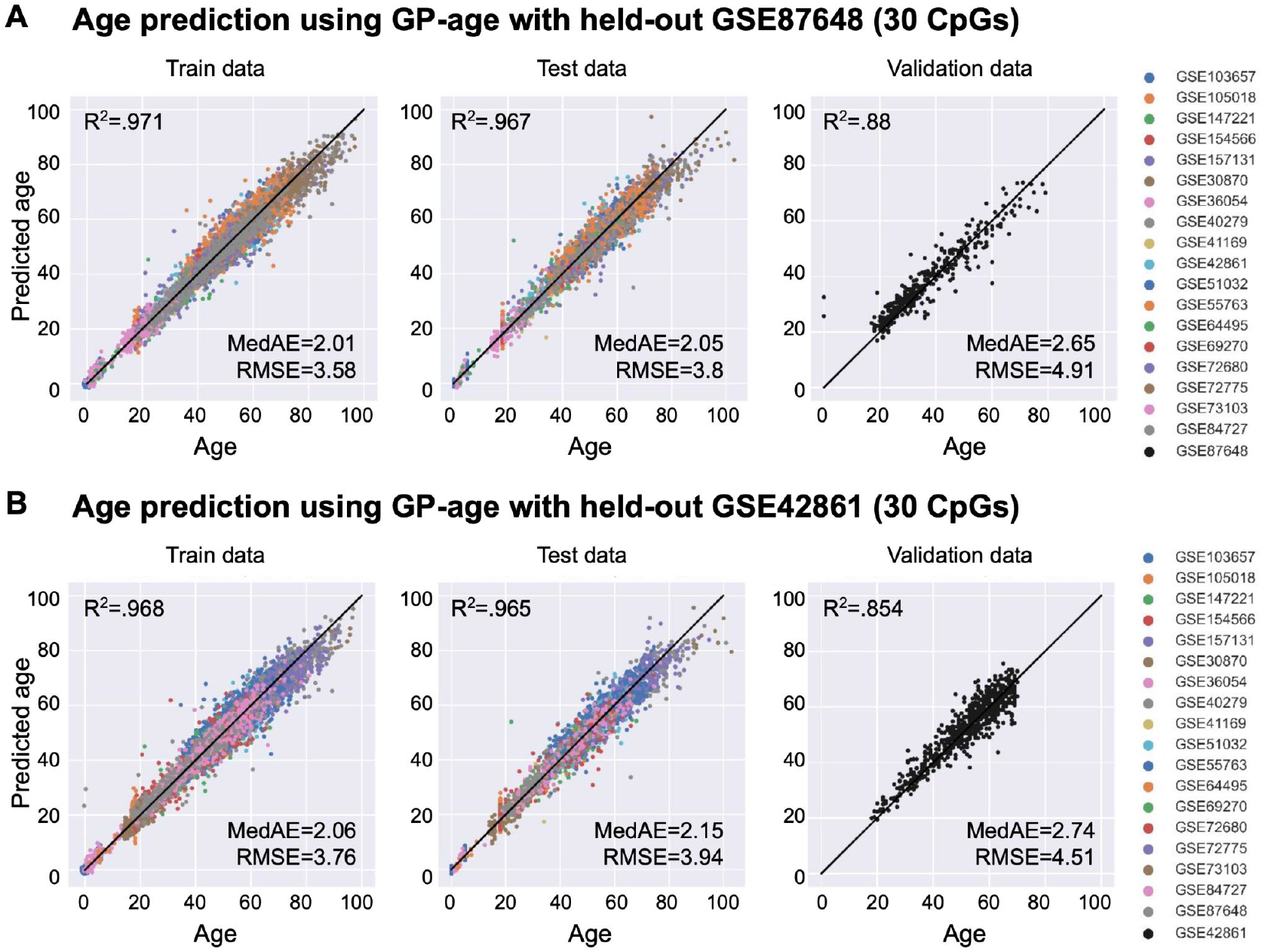
Prediction accuracy of parallel GP-age models trained with different held-out validation sets. **(A-B)** Chronological age vs. predicted age by GP-age models that were trained without the held-out validation sets GSE87648 (A) and GSE42861 (B), with 30 CpG sites, across train, test and validation set samples. Coefficient of determination R2 between prediction and age, RMSE, and MedAE are shown.

**Supplemental Fig. 3:**
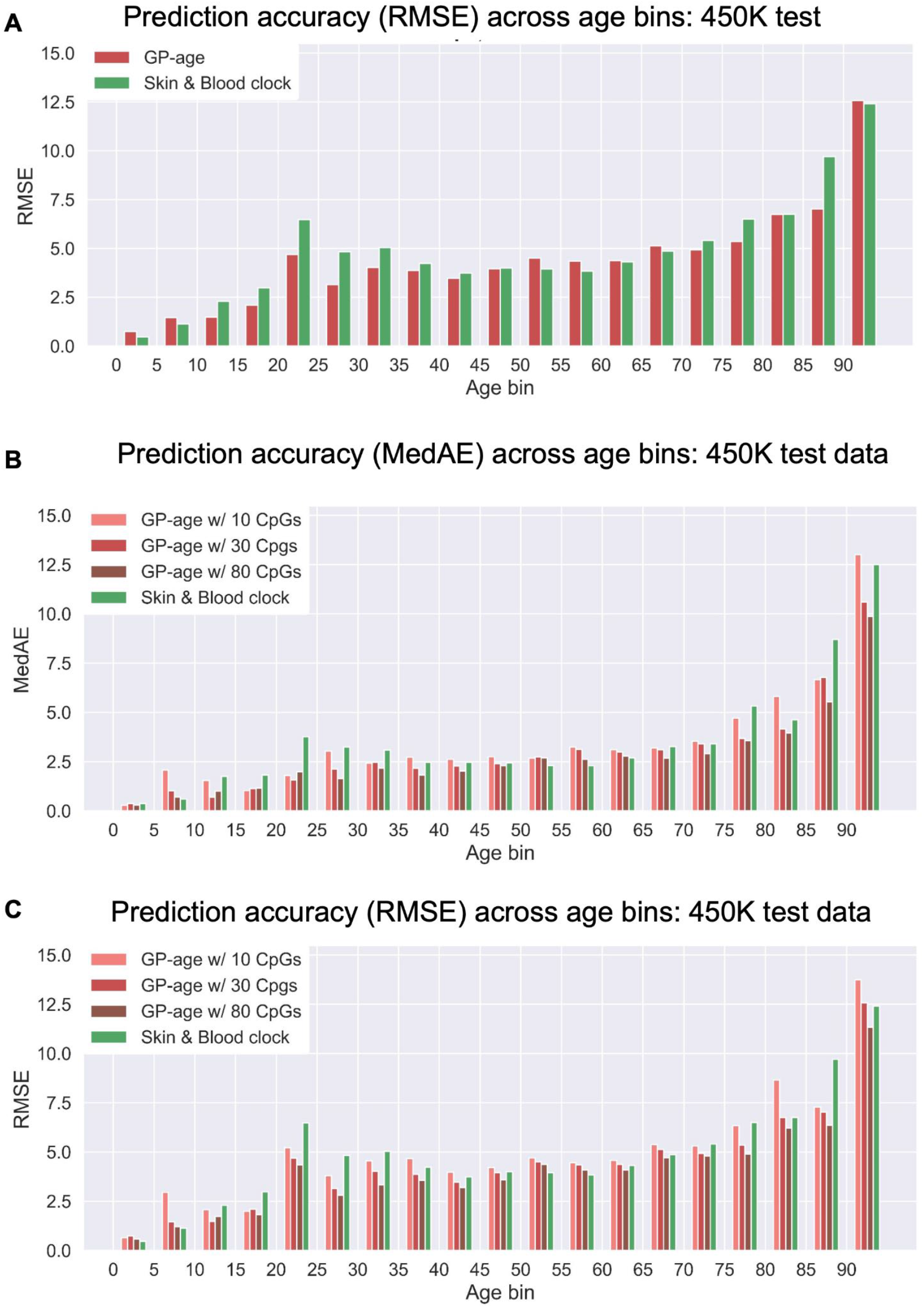
Prediction accuracy of GP-age across age bins in 450K data. **(A)** Shown are the RMSE of GP-age (with 30 CpGs) and Skin&Blood methylation clocks across different bins of donor ages. Compared to the Skin&Blood, GP-age is similarly accurate across middle-age bins, and more accurate across older- and younger-aged bins. **(B)-(C)** The MedAE (B) and RMSE (C) of GP-age of 10, 30 and 80 CpG sites and the Skin&Blood methylation clocks across different bins of donor ages.

**Supplemental Fig. 4:**
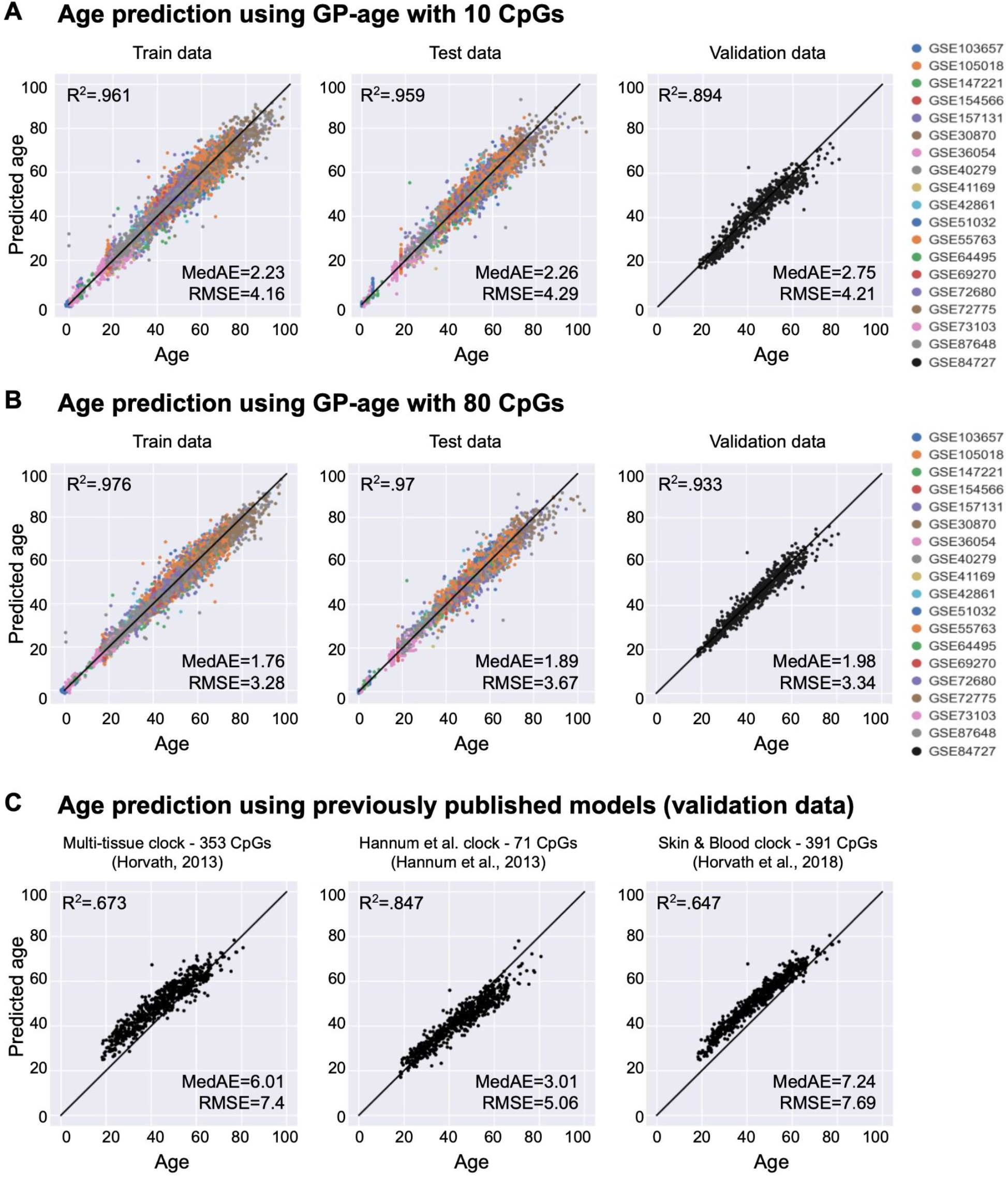
Prediction accuracy of GP-age and other clocks. **(A-B)** Chronological age vs. predicted age by GP-age models that were trained without the validation set with 10 (A) and 80 (B) CpG sites, across train, test and validation set samples. Coefficient of determination R2 between prediction and age, RMSE, and MedAE are shown. **(C)** Predictions of previously published models on the independent validation set.

**Supplemental Fig. 5:**
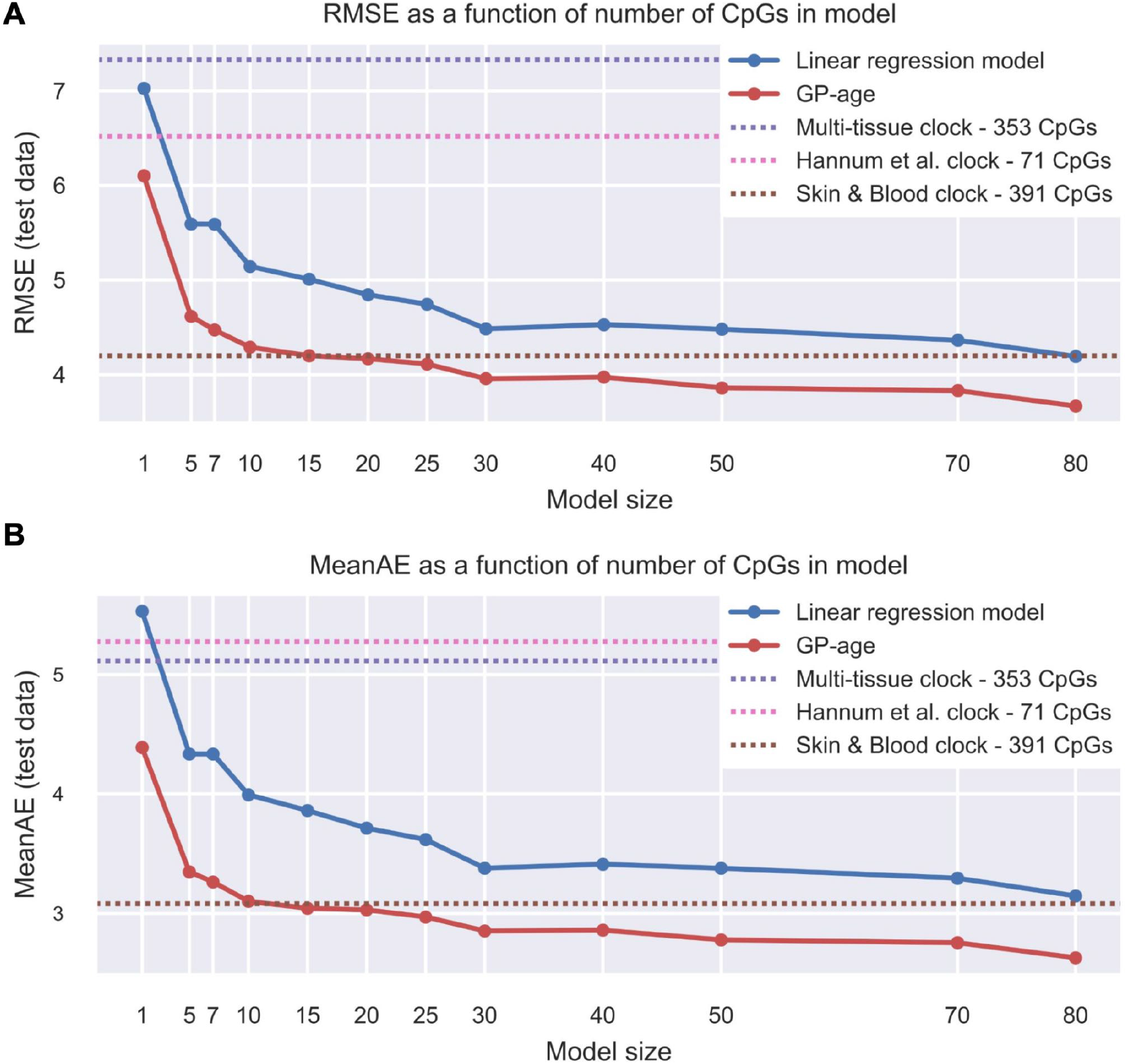
Prediction accuracy of GP-age of different model sizes. The RMSE (A) and the mean absolute error (B) of GP-age and linear model across different model sizes. GP-age outperforms previously published models. Increasing the model size increases the accuracy.

**Supplemental Fig. 6:**
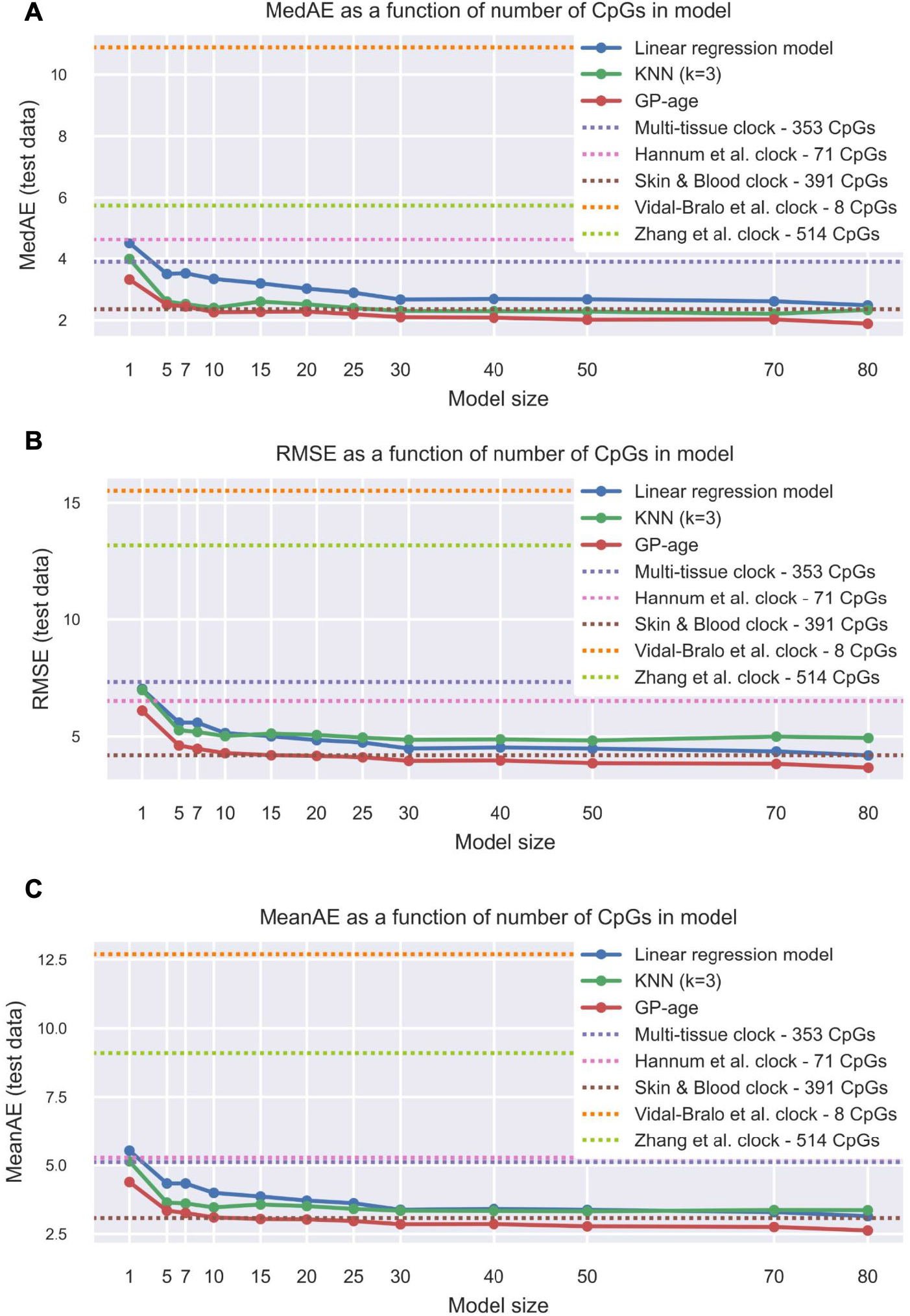
Prediction accuracy of GP-age of different model sizes, with additional models. The median error (A), RMSE (B) and the mean absolute error (C) of GP-age, linear model and KNN model across different model sizes. GP-age also outperforms the additional previously published models. Increasing the model size increases the accuracy.

**Supplemental Fig. 7:**
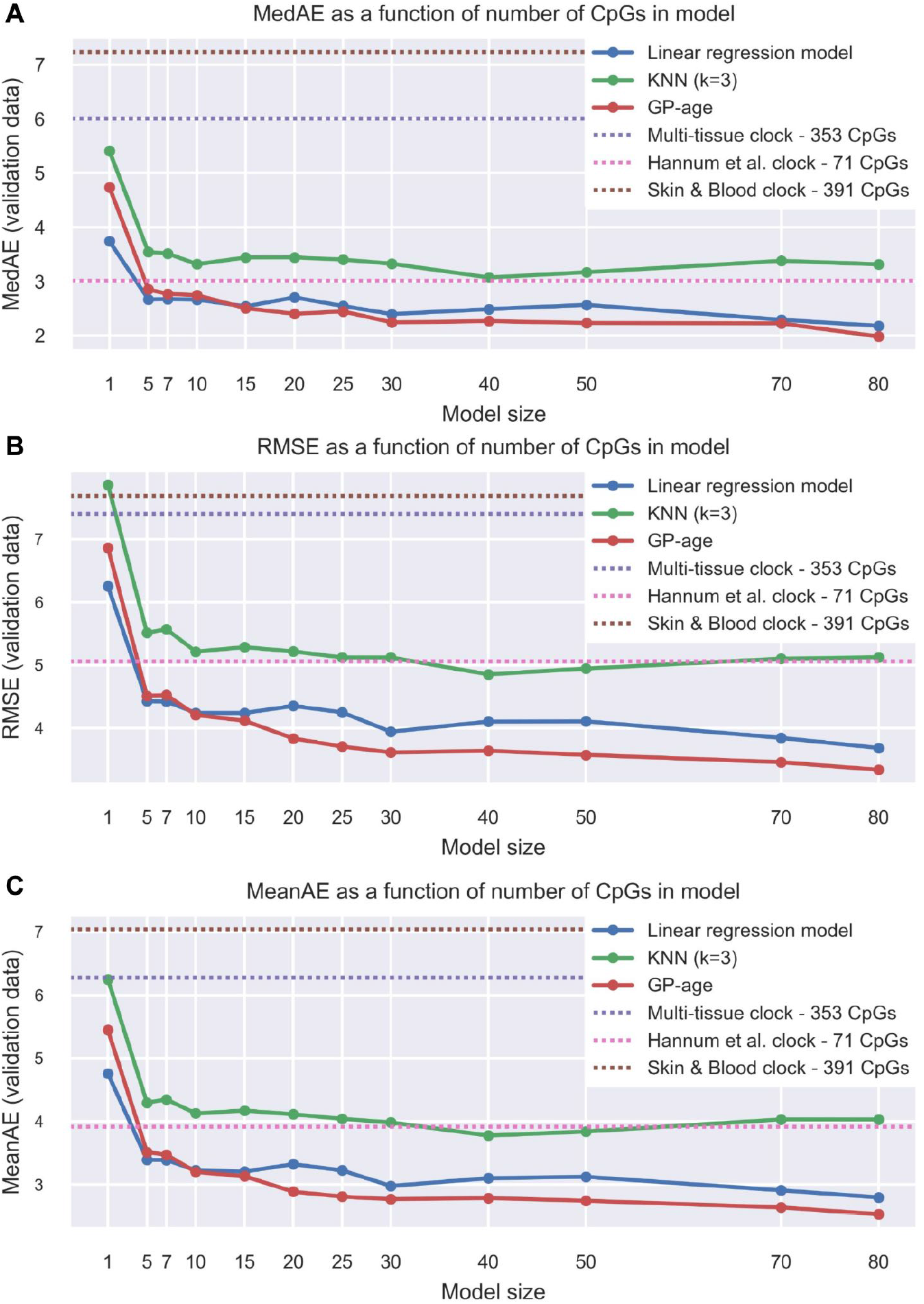
Prediction accuracy of GP-age on independent validation data. The median absolute error (A), RMSE (B), and the mean absolute error (C) of GP-age, linear model and KNN model across different model sizes, on the independent validation cohort GSE84727. GP-age outperforms previously published models. Increasing the model size increases the accuracy.

**Supplemental Fig. 8:**
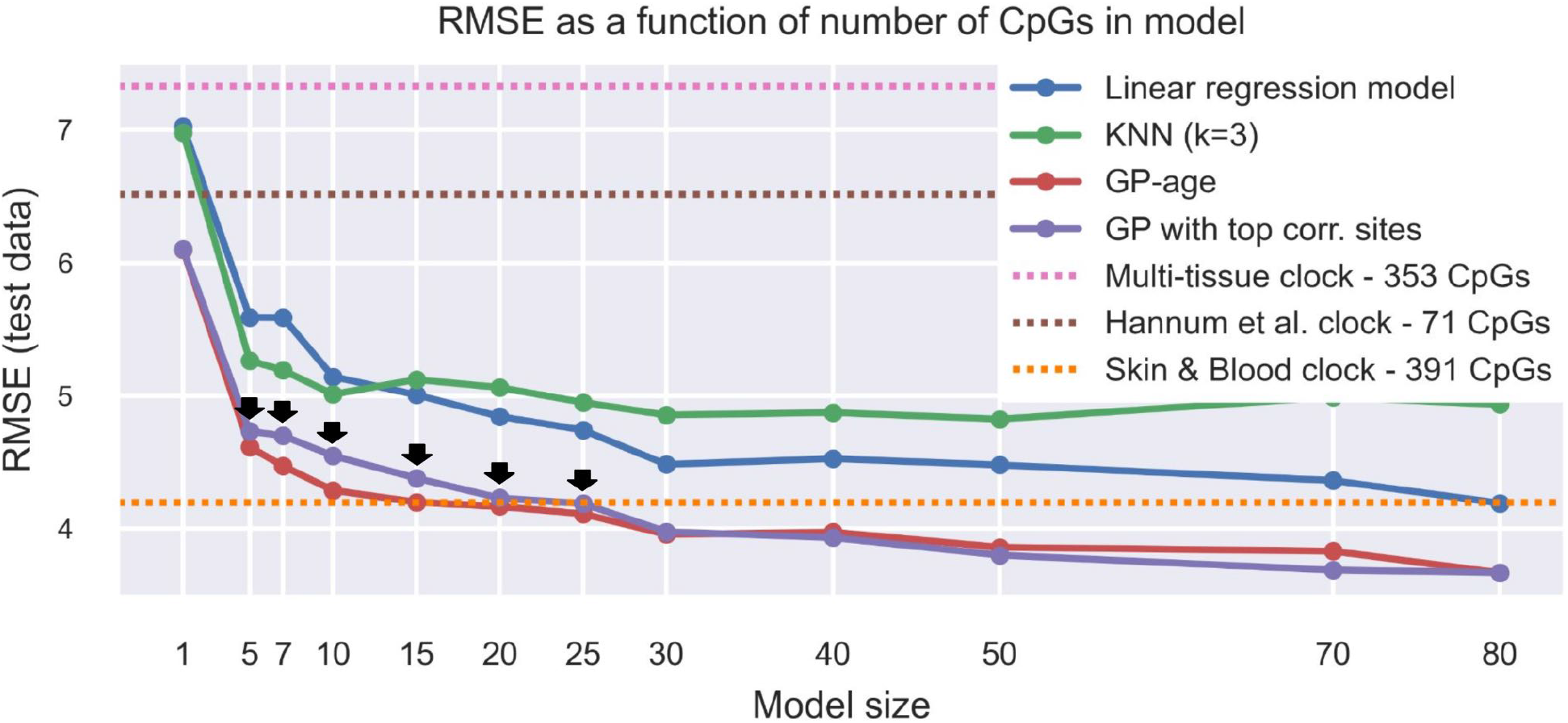
Spectral clustering yields a superior set of CpG sites for age prediction. Shown is the root mean squared error of GP-age, linear model, KNN model and a GPR model which doesn’t use spectral clustering for feature selection across different model sizes. GP-age outperforms the GPR model which uses the most correlative CpG sites for each model size when 5-25 CpG sites are used.

**Supplemental Fig. 9:**
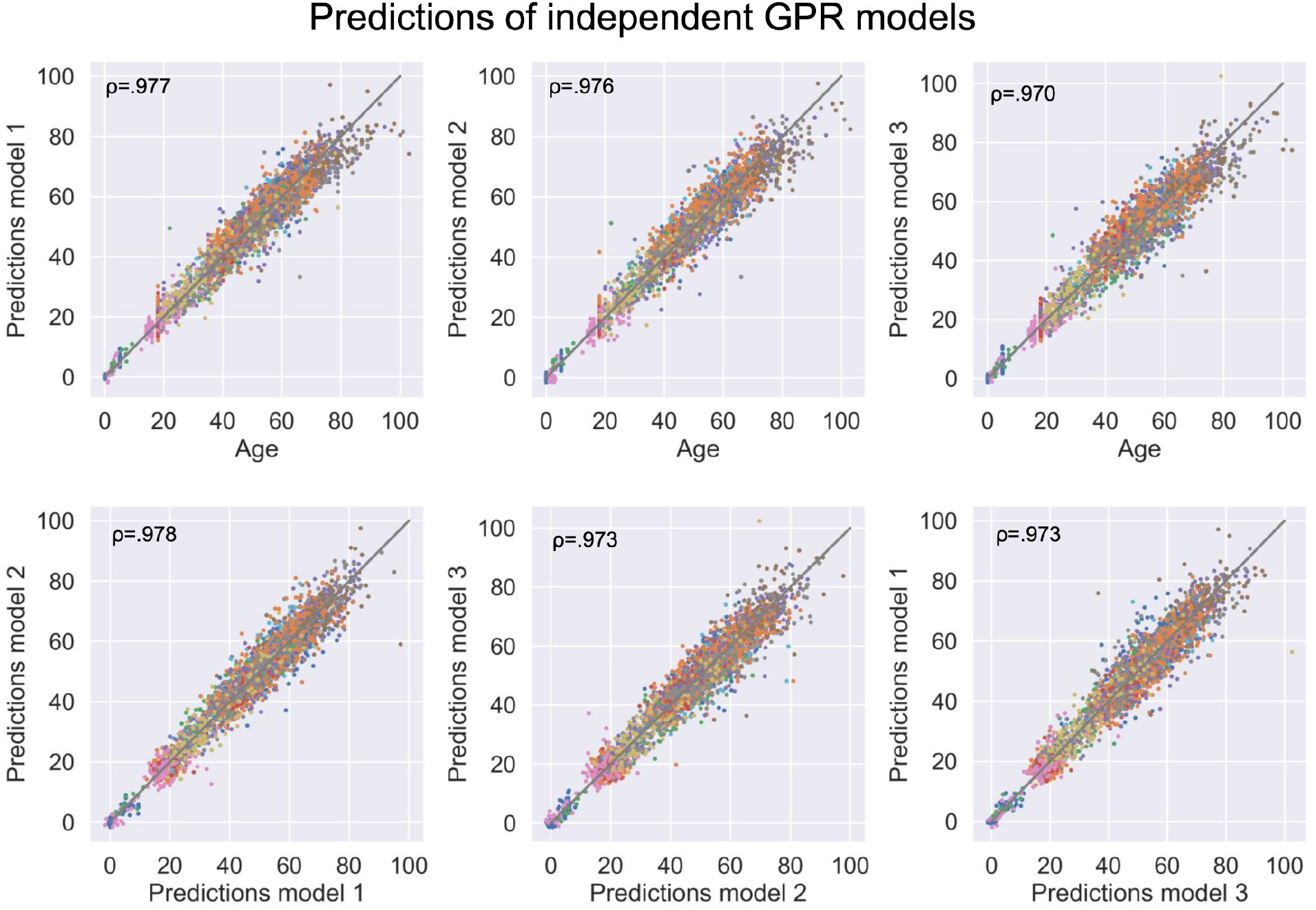
Comparison of three independent clocks. Top: shown are comparisons between chronological (x-axis) and predicted age (y-axis) in test samples, for each of the three methylation clocks. Bottom: comparisons between age predictions in each pair of models.

**Supplemental Fig. 10:**
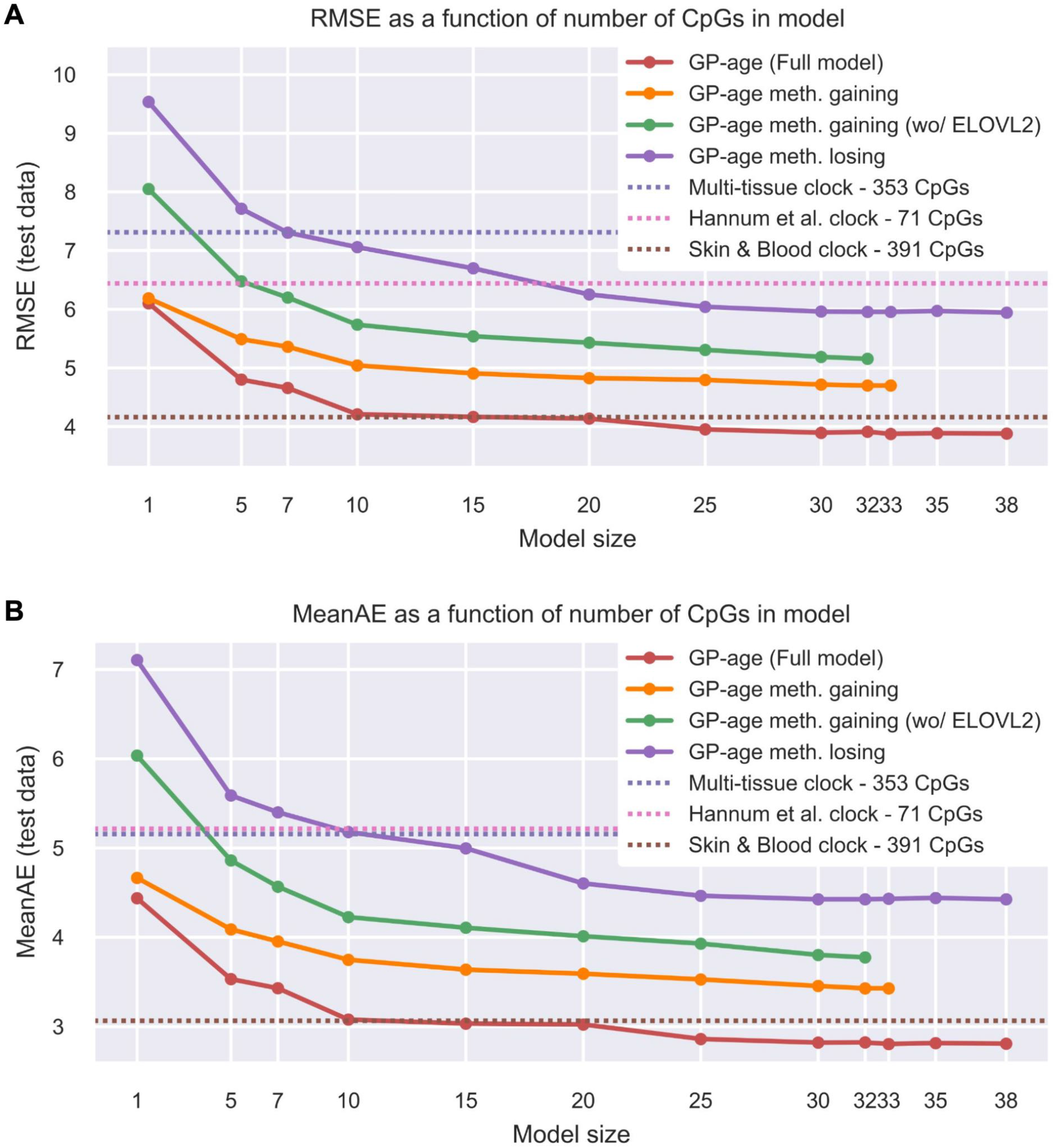
Methylation-gaining CpGs are more correlated with age than methylation-losing CpGs. RMSE (A) and mean average error (B) of GPR models. A model trained with only methylation-gaining CpG sites (orange) is more accurate than models trained with methylation-losing CpG sites (purple). This is still observed when the ELOVL2 CpG is excluded (orange). The full GP-age model outperforms these models (red).

**Supplemental Fig. 11:**
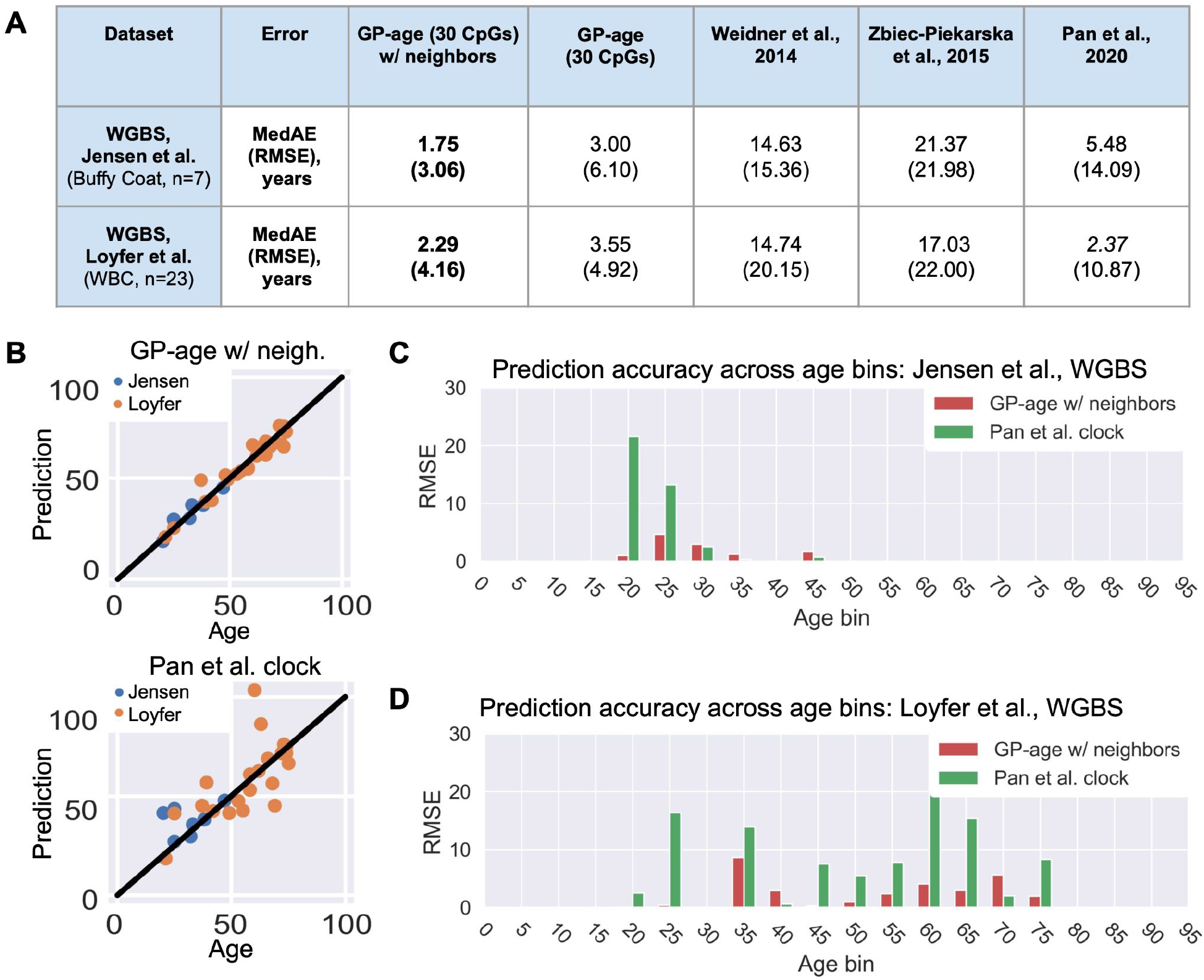
Predictive accuracy of GP-age in WGBS data. **(A)** RMSE and MedAE errors of GP-age (with and without neighbors) and reference models, across datasets. **(B)** Age prediction across the two sequencing datasets. Top: Using GP-age with a Laplace kernel for a methylation level estimation by a weighted average of methylation levels of neighboring CpGs. Bottom: Using Pan et al. clock^47^. **(C-D)** Shown are the RMSE of GP-age and Pan et al.^47^methylation clocks across different bins of donor ages for data from Jensen et al. (C) and Loyfer et al. (D). GP-age is more accurate across most age bins.

**Supplemental Fig. 12:**
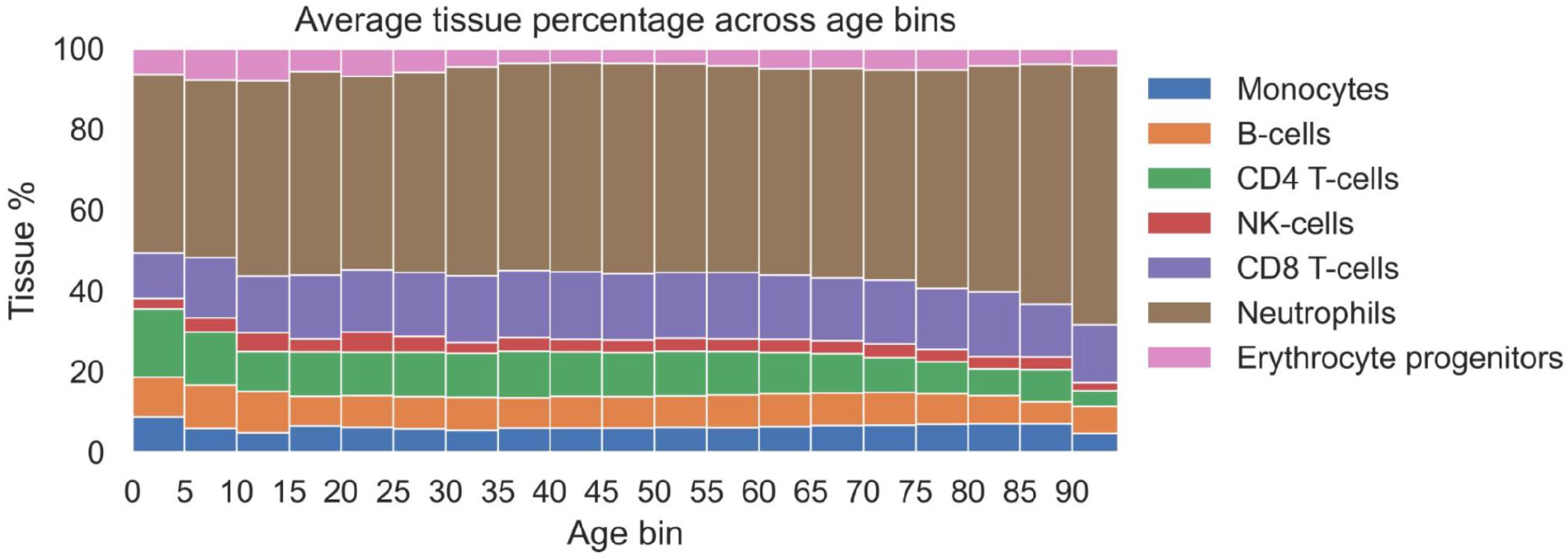
Blood cells composition across age. DNA-methylation deconvolution blood samples estimate the cellular composition of blood across different ages. While some cell types (T cells, Neutrophils) slightly change their relative proportion in the blood, these changes are far too small to explain the large differences observed in the methylation levels of age-related CpG sites (absolute change of 20-50%).

**Supplemental Table 1.**
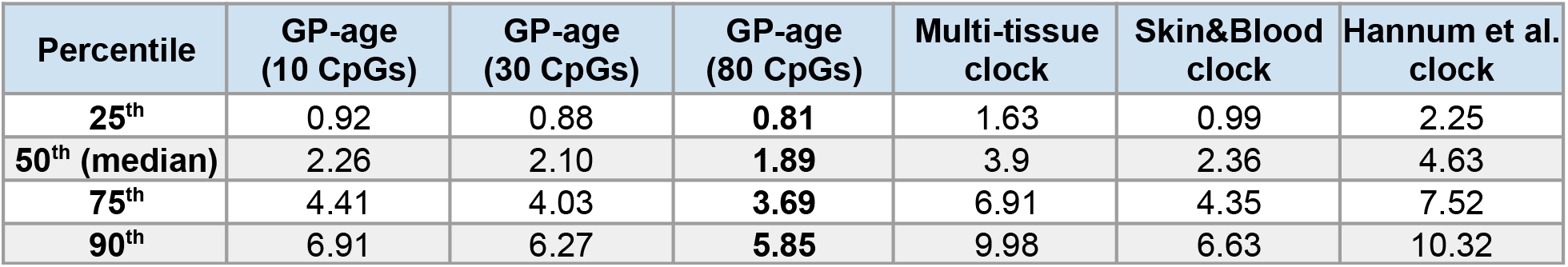
Absolute prediction errors at the 25^th^, 50^th^ (median), 75^th^ and 90^th^ percentiles of samples, for each model.

**Supplemental Table 2.** List of 1,034 age-correlative CpG sites. Sites are sorted by their absolute value Spearman correlation with age, and their correlation p-value, correlation FDR-corrected p-value and methylation range are listed.

**Supplemental Table 3.** List of 71 CpG sites with a Spearman correlation and methylation range over the defined ratios. Sites are sorted by their absolute Spearman correlation coefficient. Also shown are methylation range and neighboring genes, as well as the cluster number and whether the CpG site was included in the final list of 30 “age set” CpG sites. Last column lists, for each CpG, in which of three independent clocks it was included.

| CpG | chr | pos | Spearman<br>$\rho$ | meth.<br>range | neigh. gene | CpG type | cluster # | in final<br>set? | in clock<br>a/b/c |
| --- | --- | --- | --- | --- | --- | --- | --- | --- | --- |
| cg16867657 | chr6 | 11044877 | 0.883 | 0.345 | ELOVL2 | TSS1500 | 8 | TRUE | b |
| cg22454769 | chr2 | 106015767 | 0.809 | 0.434 | FHL2 | TSS200 | 9 | TRUE | c |
| cg06639320 | chr2 | 106015739 | 0.771 | 0.287 | FHL2 | TSS200 | 2 | TRUE | c |
| cg04875128 | chr15 | 31775895 | 0.757 | 0.361 | OTUD7A | Body | 29 | TRUE | a |
| cg19283806 | chr18 | 66389420 | -0.746 | 0.265 | CCDC102B | 5'UTR | 7 | TRUE | b |
| cg24724428 | chr6 | 11044888 | 0.737 | 0.28 | ELOVL2 | TSS1500 | 14 | TRUE | b |
| cg07553761 | chr3 | 160167977 | 0.715 | 0.236 | TRIM59 | TSS1500 | 25 | TRUE | c |
| cg24079702 | chr2 | 106015771 | 0.712 | 0.362 | FHL2 | TSS200 | 4 | TRUE | c |
| cg14556683 | chr19 | 15342982 | 0.654 | 0.215 | EPHX3 | Body | 4 | FALSE | a |
| cg07547549 | chr20 | 44658225 | 0.636 | 0.205 | SLC12A5 | Body | 2 | FALSE | b |
| cg08128734 | chr1 | 206685423 | -0.628 | 0.36 | RASSF5 | Body | 1 | TRUE | c |
| cg23500537 | chr5 | 140419819 | 0.622 | 0.223 |  |  | 2 | FALSE | a |
| cg12934382 | chr3 | 51741135 | 0.619 | 0.26 | GRM2 | 1stExon | 13 | TRUE | c |
| cg08468401 | chr3 | 14303131 | -0.613 | 0.26 |  |  | 10 | TRUE | c |
| cg05404236 | chr13 | 110437093 | 0.612 | 0.292 | IRS2 | 1stExon | 4 | FALSE | b |
| cg20816447 | chr4 | 15480781 | -0.609 | 0.221 | CC2D2A | Body | 23 | TRUE | c |
| cg17110586 | chr19 | 36454623 | 0.601 | 0.233 |  |  | 4 | FALSE | a |
| cg24466241 | chr1 | 53308908 | 0.598 | 0.223 | ZYG11A | Body | 14 | FALSE | c |
| cg18473521 | chr12 | 54448265 | 0.572 | 0.251 | HOXC4 | Body | 9 | FALSE | a |
| cg00573770 | chr2 | 145278485 | -0.564 | 0.269 | ZEB2 | TSS1500 | 5 | TRUE | c |
| cg06335143 | chr1 | 53308654 | 0.561 | 0.203 | ZYG11A | Body | 22 | TRUE | c |
| cg06155229 | chr7 | 102937535 | -0.558 | 0.221 | PMPCB | TSS1500 | 3 | TRUE | b |
| cg03032497 | chr14 | 61108227 | 0.552 | 0.273 |  |  | 30 | TRUE | c |
| cg12899747 | chr3 | 25391527 | -0.542 | 0.235 |  |  | 10 | FALSE | c |
| cg06619077 | chr1 | 47656003 | -0.541 | 0.205 | PDZK1IP1 | TSS1500 | 20 | TRUE | c |
| cg17804348 | chr1 | 3649473 | 0.539 | 0.206 | TP73 | Body | 15 | TRUE | c |
| cg00329615 | chr3 | 118706648 | -0.538 | 0.27 | IGSF11 | Body | 27 | TRUE | c |
| cg23479922 | chr5 | 16179633 | 0.533 | 0.268 | March11 | 1stExon | 17 | TRUE | a |
| cg09017434 | chr5 | 16179660 | 0.532 | 0.214 | March11 | 1stExon | 17 | FALSE | a |
| cg10501210 | chr1 | 207997020 | -0.53 | 0.57 |  |  | 11 | TRUE | c |
| cg19991948 | chr10 | 121335173 | -0.528 | 0.221 | TIAL1 | 3'UTR | 24 | TRUE | a |
| cg03738025 | chr6 | 105388694 | 0.528 | 0.216 |  |  | 14 | FALSE | b |
| cg27312979 | chr10 | 97094453 | -0.52 | 0.26 | SORBS1 | Body | 18 | TRUE | a |
| cg14766700 | chr11 | 58911067 | -0.519 | 0.21 | FAM111A | 5'UTR | 7 | FALSE | b |
| cg23186333 | chr11 | 35161900 | -0.515 | 0.232 | CD44 | Body | 6 | TRUE | b |
| cg21184711 | chr7 | 122488330 | 0.514 | 0.215 | CADPS2 | Body | 9 | FALSE | b |
| cg22730004 | chr1 | 158656718 | -0.512 | 0.211 | SPTA1 | TSS1500 | 23 | FALSE | c |
| cg19421125 | chr12 | 6882856 | -0.499 | 0.212 | LAG3 | Body | 7 | FALSE | a |
| cg15894389 | chr13 | 47470857 | -0.498 | 0.256 | HTR2A | 1stExon | 6 | FALSE | b |
| cg10835286 | chr2 | 66657913 | -0.495 | 0.247 |  |  | 3 | FALSE | c |
| cg25413977 | chr2 | 66651619 | -0.492 | 0.235 |  |  | 12 | TRUE | c |
| cg11807280 | chr2 | 66654644 | -0.49 | 0.372 |  |  | 12 | FALSE | c |
| cg22078805 | chr17 | 42432046 | 0.488 | 0.228 | FAM171A2 | Body | 26 | TRUE | c |
| cg02872426 | chr6 | 110736772 | -0.484 | 0.373 | DDO | TSS200 | 3 | FALSE | b |
| cg00303541 | chr3 | 51741280 | 0.481 | 0.204 | GRM2 | 5'UTR | 13 | FALSE | c |
| cg04295144 | chr19 | 10407184 | 0.474 | 0.239 | ICAM5 | Body | 8 | FALSE | a |
| cg19729744 | chr3 | 194752020 | -0.471 | 0.284 |  |  | 1 | FALSE | c |
| cg24794228 | chr19 | 52391166 | 0.461 | 0.214 | ZNF577 | Body | 25 | FALSE | a |
| cg17621438 | chr5 | 63461216 | -0.459 | 0.215 | RNF180 | TSS1500 | 19 | TRUE | a |
| cg05017994 | chr5 | 964562 | 0.451 | 0.21 |  |  | 3 | FALSE | a |
| cg21878650 | chr5 | 64558623 | -0.45 | 0.223 | ADAMTS6 | Body | 21 | TRUE | a |
| cg15804973 | chr6 | 137114513 | -0.449 | 0.236 | MAP3K5 | TSS1500 | 24 | FALSE | b |
| cg07797372 | chr1 | 64602654 | -0.447 | 0.223 | ROR1 | Body | 20 | FALSE | c |
| cg27152890 | chr19 | 45900241 | 0.446 | 0.248 | PPP1R13L | Body | 26 | FALSE | a |
| cg17471939 | chr1 | 3600879 | 0.439 | 0.233 | TP73 | Body | 15 | FALSE | c |
| cg18887458 | chr7 | 115995934 | -0.438 | 0.201 |  |  | 21 | FALSE | b |
| cg15243034 | chr11 | 77907656 | 0.437 | 0.235 | USP35 | Body | 2 | FALSE | b |
| cg04503319 | chr16 | 89368367 | -0.437 | 0.25 | ANKRD11 | Body | 28 | TRUE | a |
| cg03350900 | chr6 | 107955689 | 0.43 | 0.201 | SOBP | Body | 22 | FALSE | b |
| cg09809672 | chr1 | 236557682 | -0.416 | 0.223 | EDARADD | 1stExon | 16 | TRUE | c |
| cg14577707 | chr4 | 184797463 | -0.416 | 0.211 |  |  | 18 | FALSE | c |
| cg24892069 | chr10 | 33562205 | -0.412 | 0.398 | NRP1 | Body | 1 | FALSE | a |
| cg00602811 | chr2 | 145278564 | -0.409 | 0.245 | ZEB2 | TSS1500 | 27 | FALSE | c |
| cg05991454 | chr4 | 147558435 | 0.407 | 0.214 |  |  | 22 | FALSE | c |
| cg23126342 | chr13 | 67801125 | -0.406 | 0.262 | PCDH9 | Body | 18 | FALSE | b |
| cg09988805 | chr2 | 43278552 | 0.405 | 0.245 |  |  | 8 | FALSE | c |
| cg19344626 | chr19 | 16830749 | -0.403 | 0.338 | NWD1 | TSS200 | 11 | FALSE | a |
| cg20059012 | chr12 | 53613154 | -0.403 | 0.217 | RARG | Body | 23 | FALSE | a |
| cg21826784 | chr1 | 11795937 | -0.402 | 0.219 | AGTRAP | TSS1500 | 19 | FALSE | c |
| cg07164639 | chr6 | 110736958 | -0.401 | 0.245 | DDO | TSS1500 | 5 | FALSE | b |
| cg11693709 | chr15 | 40542019 | -0.4 | 0.31 | PAK6 | 5'UTR | 16 | FALSE | a |

